# Tree reconstruction guarantees from CRISPR-Cas9 lineage tracing data using Neighbor-Joining

**DOI:** 10.1101/2024.08.27.610007

**Authors:** Sebastian Prillo, Kevin An, Wilson Wu, Ivan Kristanto, Matthew G. Jones, Yun S. Song, Nir Yosef

**Affiliations:** Department of Electrical Engineering and Computer Sciences, University of California, Berkeley, CA 94720, USA; Department of Systems Immunology, Weizmann Institute of Science, Rehovot 7610001, Israel; Department of Statistics, University of California, Berkeley, CA 94720, USA; Department of Molecular and Cell Biology, University of California, Berkeley, CA 94720, USA; Center for Personal Dynamic Regulomes, Stanford University, Stanford, CA 94305, USA

## Abstract

CRISPR-Cas9 based lineage tracing technologies have enabled the reconstruction of single-cell phylogenies from transcriptional readouts. However, developing tree-reconstruction algorithms with theoretical guarantees in this setting is challenging. In this work, we derive a reconstruction algorithm with theoretical guarantees using Neighbor-Joining (NJ) on distances that are moment-matched to estimate the true tree distances. We develop a series of tools to analyze this algorithm and prove its theoretical guarantees. When the parameters of the data generating process are known and there is no missing data, our results align with established results from common evolutionary models, such as Cavender-Farris-Neyman and Jukes-Cantor. However, to account for the realistic case where the parameters of the data generating process are not known and there is missing data, we develop new theory that shows for the first time that it is still possible to obtain reconstruction guarantees in the CRISPR-Cas9 case and in other models of evolution. Empirically, we show on both simulated lineage tracing data and on real data from a mouse model of lung cancer the improved performance of our method as compared to the traditional use of NJ.

## 1 Introduction

A foundational principle of biology is that cells continuously divide to give rise to tissues and organisms. This principle has been useful in understanding dynamic processes across biology, from normal development to the generation of tumors and seeding of metastatic lesions. Phylogenetic trees are useful models for studying these processes as they can describe the growth process from founding cells to current-day cell populations. Recent developments in CRISPR-Cas9 based lineage tracing technologies have enabled the reconstruction of these ‘historical’ single-cell phylogenies from only present-day cell populations. Generally, these technologies work by stochastically introducing heritable mutations in the form of insertions and deletions (“indels”) at a defined locus (e.g., the 3’ UTR of a fluorescent protein) that can be subsequently read out with single-cell genomic technologies like single-cell RNA-seq (scRNA-seq) [1, 2, 3, 4, 5, 6, 7]. The accumulated edits introduced by the CRISPR-Cas9 machinery can then be used to infer clonal relationships, or phylogenetic trees describing the emergence of cell populations. Thus far, these technologies have enabled a deeper understanding of embryonic development and tumor growth in animal models [8, 9, 10, 11, 12].

Despite the rapid technological developments enabling CRISPR-Cas9 lineage tracing, a key challenge is the computational reconstruction of cellular phylogenies. Many algorithms have been proposed to reconstruct the single-cell phylogeny’s topology from these CRISPR-Cas9 mutations [13, 14, 15, 16, 17, 18, 19]. However, for few of these algorithms do we have any guarantees that the reconstructed topology will be accurate, even if we assume that there are infinitely many mutation sites evolving under the CRISPR-Cas9 model. The challenges that underlie the CRISPR-Cas9 case as compared to prevalent case studies that rely on classical models such as Cavender-Farris-Neyman (CFN) and Jukes-Cantor (JC) are (1) the presence of missing data and (2) unknown model parameters such as mutation rate and transition probabilities. Recent work from our group [19] proposed character-based algorithms with theoretical guarantees for the CRISPR-Cas9 setting. However, distance-based methods such as Neighbor-Joining (NJ) [20] - which work well in many settings [21, 22] and in practice for CRISPR-Cas9 data [15, 16] - have no formulated guarantees for CRISPR-Cas9 data yet [19].

In this work, we build upon commonly used methods in statistical phylogenetics to propose a distance-based algorithm with theoretical reconstruction guarantees from CRISPR-Cas9 data. Instead of applying NJ to Hamming distances (as is usually done), we apply NJ on *corrected* distances which are estimates of the *true pairwise tree distances*. This is a standard approach in statistical phylogenetics which is routinely applied to models such as CFN and JC [23, 24, 25]. The main challenge is to prove theoretically that it works for the CRISPR-Cas9 setting. Our first main contribution is therefore to formalize this approach in a generic manner, prove its guarantees for the general case (handling any evolutionary model), and then show its correctness for the case of CRISPR-Cas9. Other major difficulties arise due to the presence of missing data and because the parameters of the model such as the mutation rate and the state probabilities are not known. Our second main contribution is a general formulation of this setting, with provable guarantees that depend on one’s ability to infer the parameters of the model. We demonstrate how this formulation applies to the CRISPR-Cas9 setting by presenting an algorithm that couples approximation of the missing parameters with tree inference.

To analyze our algorithms and prove our bounds, we develop a set of elementary analytical tools that allow us to estimate how errors propagate throughout the algorithm. A key quantity that arises in these estimates and underlies our theory and bounds is the *minimum increment functional*, which measures how flat a function is locally, in the worst case. Our general theorems include as a corollary known results for the CFN and JC models. The bounds we derive are asymptotically similar to those derived recently by our lab [19] for character-based methods, providing further support to the results presented there about the relative importance of the different experimental parameters in CRISPR-Cas9 lineage tracing. For example, the Cas9 cutting rate is found to be a more critical parameter than the state diversity, and similarly, increasing the number of characters is found to be more important than increasing the state diversity. Our theorems are general and can be used to derive algorithms with theoretical guarantees for other models beyond the CRISPR-Cas9 setting, notably - to reiterate - in cases when there is missing data and model parameters are not known.

Empirically, we show there is consistently increased performance of our method as compared to NJ applied to raw Hamming distances for the CRISPR-Cas9 model, both on simulated data as well as on real data from a mouse model of lung adenocarcinoma [11].

In the sections that follow, we first provide definitions and key results from prior work. Then, we present our theoretical framework and novel technical results, which apply not only to the CRISPR-Cas9 model, but also to classical models such as CFN and JC. Finally, we show empirically, on simulated and real data, the improved performance of our method as compared to the usual application of NJ to raw Hamming distances. Together, we demonstrate that best practices from statistical phylogenetics carry on to the CRISPR-Cas9 lineage tracing setting, and we provide a general theoretical justification to that approach.

## 2 Problem Setup

This work is written in the language of statistical models to make our general results precise. We start by defining what we mean by an ‘evolutionary model’ along with useful definitions and prior work before introducing the CRISPR-Cas9 model (which is a particular case of an evolutionary model):

### Definition 1

(Evolutionary model). An *evolutionary model* is a statistical model

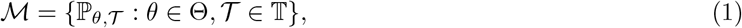

where Θ parameterizes a continuous-time Markov chain (CTMC) with state space 𝕊 (for example, 𝕊 = {*A, C, G, T*} for the Jukes-Cantor model), 𝕋 is a set of edge-weighted rooted binary trees, and ℙ_*θ,T*_ is the probability measure for the stochastic process resulting from running the CTMC parameterized by *θ* down the tree 𝒯 ; importantly, at each internal node, the chain is copied and continues to evolve independently over each child branch. For a positive integer *k*, we denote ℳ_*k*_ the statistical model resulting from *k* i.i.d. realizations of the model 𝒫, in other words:

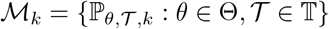

We say that ℳ_*k*_ is an *evolutionary model with k similarly evolving characters*. The value of this stochastic process at node *v* for character *i* is denoted by 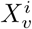. This way, *X* is a random matrix indexed by the vertices of 𝒯 (including the internal nodes) and 1 ≤ *i* ≤ *k* with entries in 𝕊. For ease of notation, we will drop the subscript *k* when writing ℙ_*θ,T, k*_. Expectations with respect to ℙ_*θ,T, k*_ will be written as 𝔼_*θ,T, k*_ and similarly we will drop the subscript *k*.

In what follows, we let *V* (𝒯) denote the set of vertices of 𝒯, *L*(𝒯) the set of leaves of 𝒯, and *r*(𝒯) the root of 𝒯. It is important to clarify that in this work, when we refer to a rooted binary tree, we consider the root to have degree 1 (rather than 2). This is important since in a single cell phylogeny the progenitor cell does not divide immediately. Furthermore, this class of rooted binary trees is strictly more general than when we restrict the root to have degree 2 (by making the root edge have length 0). Additionally, when we talk about the *diameter* of a tree, we mean the maximum path length between any two nodes. The data we have available to reconstruct the tree is called the ‘character matrix’, formalized as follows:

### Definition 2

(Character matrix). Let M_*k*_ be an evolutionary model with *k* similarly evolving characters. The restriction *X*_*L*(𝒯)_ of *X* to the leaves of 𝒯 is called the *character matrix*, which has size |*L*(𝒯)| × *k*.

The goal of phylogenetic tree reconstruction is to use the character matrix *X*_*L*(𝒯)_ to reconstruct the topology of 𝒯. Two broad families of algorithms for solving this problem exist: *distance-based methods* and *character-based methods*. In distance-based methods, the character matrix *X*_*L*(𝒯)_ is summarized into a *dissimilarity matrix D* of size |*L*(𝒯)| × |*L*(𝒯)| which is assumed by definition to be non-negative, symmetric, and zero on the diagonal, and then a *distance-based algorithm* such as Neighbor-Joining is used to reconstruct the tree using *D*. In contrast, character-based methods operate on the character matrix *X*_*L*(𝒯)_ directly and try to optimize measures of tree fit such as parsimony or likelihood. In this work, we are concerned with developing distance-based methods with theoretical guarantees for the CRISPR-Cas9 setting. Character-based methods with guarantees have been recently derived by our lab [19].

We first formalize what we mean by a distance-based algorithm:

### Definition 3

(Dissimilarity function (or matrix)). Let *A* be a set. A *dissimilarity function (or matrix) over A* is a non-negative function *D* : *A* × *A* → ℝ_≥0_ that satisfies:

- *D*(*a*_1_, *a*_2_) = *D*(*a*_2_, *a*_1_), ∀*a*_1_, *a*_2_ ∈ *A*.
- *D*(*a, a*) = 0, ∀*a* ∈ *A*.

### Definition 4

(Unrooted distance-based tree reconstruction algorithm). An *unrooted distance-based tree reconstruction algorithm* – or uDBA for short – is a deterministic algorithm (and thus a function) 𝒜 that given a dissimilarity function *D* : *L* × *L* → ℝ_≥0_ over a set *L*, returns an unrooted tree 𝒯 with leaf set *L*.

A wealth of theoretical research exists on uDBAs such as Neighbor-Joining. Let *d*_*𝒯*_ (*u, v*) be the distance between *u* and *v* in the edge-weighted unrooted binary tree 𝒯 and let *l*_min_(𝒯) be the smallest edge length of T. Then, a result by Atteson [23] shows that if a dissimilarity function *D* satisfies |*D*(*u, v*) −*d*_*𝒯*_ (*u, v*) | *< l*_min_(𝒯)*/*2 for all leaves *u, v*, then Neighbor-Joining run on *D* will return the correct unrooted tree topology of 𝒯. Other uDBAs enjoy this property such as FastME and GreedyBME [26]. Generally, we have the following, known as the *Atteson condition* or *l*_∞_*-radius* for an uDBA:

### Definition 5

(*l*_∞_-radius, or Atteson condition). An uDBA is said to have an *l*_∞_*-radius* of *R* if whenever |*D*(*u, v*) − *d*_*𝒯*_ (*u, v*)| *< R l*_min_(𝒯) for all leaves *u, v* of a weighted unrooted binary tree 𝒯, then running the uDBA on *D* will return the correct unrooted tree topology of 𝒯.

It is known that an *l*_∞_-radius of 1*/*2 is optimal [23], and thus Neighbor-Joining has an optimal *l*_∞_-radius. Unfortunately, the Atteson condition is not well suited to dissimilarity functions *D* such as the Hamming distance because they are not linearly related to tree distance. Indeed, for a simple model such as CFN, if two leaves *u, v* are at a tree distance of *d*_*𝒯*_ (*u, v*) = *t*, then their expected binary Hamming distance for one character is 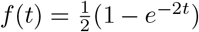. Because of this, distance-based methods such as NJ are typically applied to ‘corrected’ distances that are better estimates of the tree distance. This is done by leveraging the mapping *f*^−1^. Concretely, Neighbor-Joining is typically run on the corrected distances given by 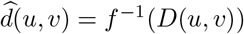. One must be careful when inverting *f* because *D*(*u, v*) might lie outside the image of *f*, which is easily fixed with *clipping*, wherein *D*(*u, v*) is capped to the maximum allowed value of *f*. Using this approach, theoretical guarantees have been derived for models such as CFN and JC [23, 24, 25]. In this work, we adapt these techniques to the CRISPR-Cas9 setting. However, this is challenged by the fact that the CRISPR-Cas9 model has (1) missing data, and (2) unknown parameters, issues which are not addressed in the classical analysis.

We now formally define the CRISPR-Cas9 evolutionary model:

### Definition 6

(CRISPR-Cas9 evolutionary model). The CRISPR-Cas9 evolutionary model ℳ_*k*_ with *k* similarly evolving characters is defined as follows:

a. The set of trees 𝕋 is the set of ultrametric weighted rooted binary trees of height exactly 1. That is to say, we assume without loss of generality that the experiment is run for exactly one unit of time.
b. The underlying CTMC starts in the state 0, called the *unmutated* or *unedited* state. A character in the 0 state mutates at a rate of *λ >* 0. These mutations are *irreversible*, meaning that once a character mutates, then it cannot mutate again. When a character in the 0 state mutates, it acquires a new state from the set 𝕊 = ℤ_+_, where state *j* is acquired with probability *q*_*j*_. Here, 𝕊 represents the set of possible indels formed by the CRISPR-Cas9 mutation process. This way, the CTMC is parameterized by *λ >* 0 and the state probabilities *q*_*j*_ ≥ 0, so that *θ* = (*λ, q*_1_, *q*_2_, …). As usual, the model is run with *k* i.i.d. characters to give ℳ_*k*_.

Note that our definition of the CRISPR-Cas9 evolutionary model does not include missing data. We will address missing data in Section 4.2.

## 3 Overview of Results

In Section 4.1, we show that when there are no missing data and the parameters of the CRISPR-Cas9 model are known, it is possible to reconstruct the tree topology with 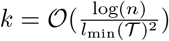 characters using the distance-correction scheme from statistical phylogenetics. For this, we first generalize the distance-correction scheme from statistical phylogenetics to any evolutionary model and analyze the number of characters *k* needed. We focus on ultrametric trees as in the case of CRISPR-Cas9. The main result is Theorem 1. A key ingredient of this theorem is Lemma 1, which controls how errors in the raw dissimilarities *D* propagate to errors in the corrected distances 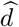.

Next, in Section 4.2, we explain how the method and proofs can be adjusted to deal with missing data. We provide a general result in Theorem 3. We specialize this theorem to the CRISPR-Cas9 setting to show that when each entry in the character matrix is missing marginally with probability *p*_missing_, it is possible to reconstruct the tree topology with 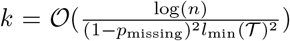 characters. This is the result of Corollary 1. Note that the dependency on 1 − *p*_missing_ in the denominator is quadratic, which is better than the cubic dependency our lab derived in [19] for character-based methods.

Furthermore, in Section 4.3, we explain how the method and proofs can be adjusted to deal with unknown model parameters, which requires new techniques and represents the biggest technical contribution of our work. We provide a general result in Theorem 4. A key ingredient is Lemma 5, which controls how errors in the model parameters – and thus errors in *f* – propagate to errors in the corrected distances 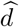. We specialize this theorem to the CRISPR-Cas9 setting to show that when neither the mutation rate *λ* nor the state probabilities *q, q*, … are known, it is still possible to reconstruct the tree topology with 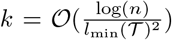 characters. This is the content of Theorem 5. It follows from applying our techniques for missing data, that when there are both missing data and unknown parameters, it is possible to reconstruct the tree topology with 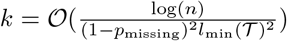 characters in the CRISPR-Cas9 setting. This is the content of Corollary 2, and completes the main theoretical contributions of our work.

Finally, we show consistently improved performance of our method on empirical data obtained from simulations as well as on real data from a mouse model of lung adenocarcinoma [11], compared to the usual application of NJ to CRISPR-Cas9 data using raw Hamming distances.

Our framework can be applied to other settings where trees are not necessarily ultrametric. In *SI Appendix, SI Text* B we provide a version of Theorem 1 that applies to evolutionary models with non-ultrametric trees and where the underlying Markov chain is stationary and reversible. This is the content of Theorem 7. With this, we derive known bounds for CFN and JC as corollaries in Corollary 3 and Corollary 4 respectively. We also provide the counterpart of Theorem 4 (which allows for unknown parameters) in Theorem 8. This demonstrates the generality of our framework.

## 4 Theoretical Results

In this section, we present the main theoretical results. All proofs are deferred to *SI Appendix, SI Text* A.

### 4.1 Analysis of distance-correction scheme with known model parameters

In the distance-correction scheme, the *raw distances D*, such as Hamming distances (or, more generally, any kind of distances such as weighted Hamming distances) are inverted to obtain corrected distances which are estimates of pairwise tree distance. These are then used in an uDBA to estimate the tree topology. Thus, it is crucial to understand how errors in the raw distances *D* translate to errors in the estimated pairwise tree distances 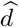. Bounding these errors would allow us to derive theoretical results via the Atteson condition [23].

The expected raw distance function *f*(*t*) is the key object in this scheme; defining clip_*a*_(*b*) = min{*a, b*}, the corrected distances are given by 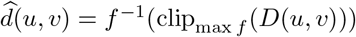. For example, for the CFN model with a fixed mutation rate of 1.0 and where the Hamming distance is normalized by the number of characters (such that it lies between 0 and 1) we have 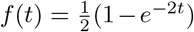 and therefore 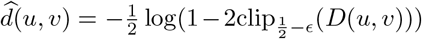 where *ϵ >* 0 is a user-chosen parameter that is used to ensure proper clipping (since *f*(*t*) ≥ 1*/*2 is not attainable). Intuitively, it is harder to invert *f* at points where *f* is ‘flat’, in other words, at points where the gradient of *f* is close to 0. Indeed, in this case small errors in the observed raw distance *y* translate to large errors in our estimate of the true distance *t*, as illustrated in Figure 1. Lemma 1 makes this rigorous and explains what error in *y* is sufficient to obtain a small error in *t*. To introduce the lemma, we first define the *minimum increment functional* Δ, which measures how flat the function *f* is locally, in the worst case:

**Figure 1:**
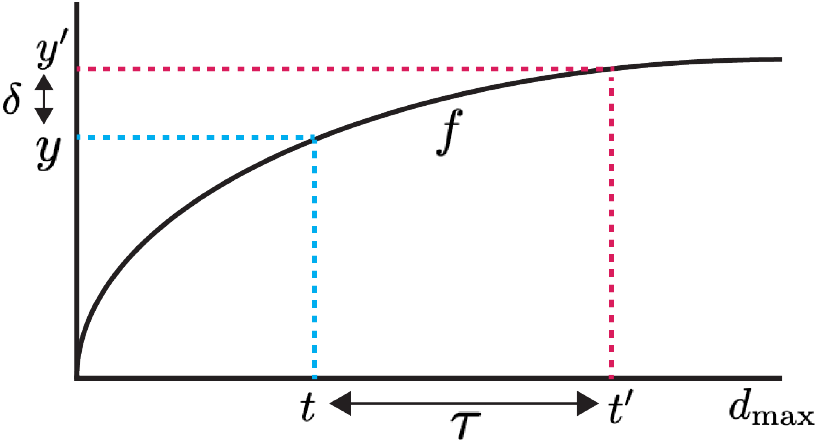
In the distance-correction scheme, the observed raw distance *y* is inverted through the expected raw distance function *f* : [0, *d*_max_] → ℝ_≥0_ to get an estimate of the true pairwise tree distance *t*. Because *y* is noisy, small errors *δ* in *y* may translate to large errors in *t*, as illustrated in the figure. The ‘flatter’ *f* is at the point *f*(*t*), the larger the error. Lemma 1 formalizes this intuition and explains how large the error *δ* in *y* can be to achieve a desired maximum error of *τ* in *t*. To be precise, for a given *τ*, Lemma 1 shows that a universal value of *δ* which works for any *t* in the interval [0, *d*_max_] is 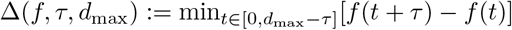.

#### Definition 7

(Minimum increment functional Δ). Let 0 *< a* ≤ *d*_max_ be real numbers. Let *f* : [0, *d*_max_] → ℝ_≥0_ be a continuous, strictly increasing function, and let *τ* ∈ [0, *a*]. We define the *minimum increment functional* Δ as:

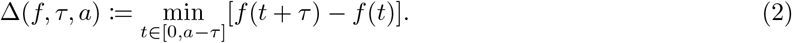

Intuitively, the minimum increment Δ(*f, τ, a*) is a (tight) lower bound on the increment that *f* attains when evaluating it on an input that is larger by *τ*, all while restricting the evaluations of *f* to the interval [0, *a*]. Thus, if *f* looks ‘flat’ in some window of length *τ*, it will have a small minimum increment.

With this, we can state our first key lemma:

#### Lemma 1

(Minimum increment lemma). Let *d*_max_ *>* 0. Let *f* : [0, *d*_max_] → ℝ_≥0_ be a continuous, strictly increasing function. Let *t* ∈ [0, *d*_max_] and *y* = *f*(*t*). Let *y*^*′*^ ≥ 0 and *τ* ∈ [0, *d*_max_]. Then, we have:

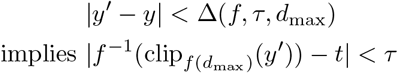

In fact, the following stronger statement holds:

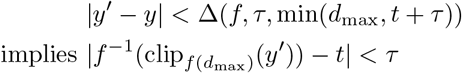

In the particular case of the Atteson condition, where we want to achieve an error at most *Rl*_min_(𝒯) for all estimated pairwise tree distances, Lemma 1 tells us that it suffices to have an error at most Δ(*f, Rl*_min_(𝒯), *d*_max_) in the raw distances *D*. We formalize this in the following proposition, tailored to the CRISPR-Cas9 setting where the trees are ultrametric:

#### Proposition 1

(Accurate ultrametric rooted tree reconstruction assuming pairwise distances can be reasonably estimated from *f*). Suppose that 𝒯 is an ultrametric weighted rooted binary tree with height *h >* 0. Suppose that *D* is a dissimilarity matrix over the leaves of 𝒯, and *f* : [0, 2*h*] → ℝ_≥0_ is a continuous strictly increasing function with *f*(0) = 0. If is an uDBA with *l*_∞_-radius *R*, and if for every pair (*u, v*) of leaves we have that

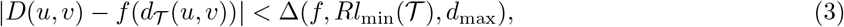

then, if we define the random dissimilarity matrix 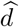 over *L*(𝒯) ∪ {*r*(𝒯)} as

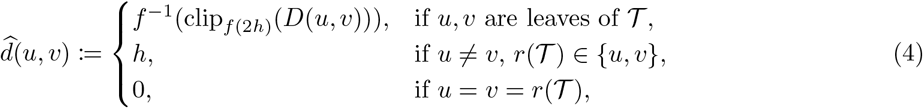

we have that running 𝒜 on *d* and 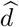 rooting the resulting tree at *r*(𝒯) gives the correct rooted tree topology of 𝒯.

We are almost ready to state our first general theorem, which applies to evolutionary models with ultrametric trees, as in the CRISPR-Cas9 setting. We just require the following definitions. The first specifies what kind of dissimilarity matrices our method applies to. Informally, these need to be averages over the *k* characters:

#### Definition 8

(Dissimilarity matrix associated to an evolutionary model and dissimilarity function). Let ℳ_*k*_ be an evolutionary model with *k* similarly evolving characters with state space 𝕊 and let *D* be a dissimilarity function over 𝕊. Over *V* (𝒯) we define the random dissimilarity matrix *D*_*k*_ : *V* (𝒯) × *V* (𝒯) → ℝ_≥0_ as follows:

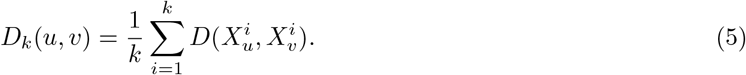

We call *D*_*k*_ the *dissimilarity matrix associated to ℳ*_*k*_ *and D*.

For example, the (average) Hamming distance is the dissimilarity matrix *D*_*k*_ associated to any evolutionary model ℳ_*k*_ and the indicator for inequality *D*(*x, y*) = 𝟙{*x* ≠ *y}*. Other dissimilarities *D*_*k*_ such as the weighted Hamming distance arise from other choices of *D*.

Next, it is trivial but important to note that the marginal distribution of the characters of two leaves in a tree 𝒯 depends only on the subtree they induce, which we call a 𝒴-tree since it is shaped like an inverted ‘Y’:

#### Definition 9

(𝒴-tree). Let *h, t >* 0 be real numbers with *t* ≤ 2*h*. We define 𝒴 (*t, h*) to be an ultrametric weighted rooted binary tree of height *h* with exactly two leaves labelled 1 and 2, separated by a tree distance of *t*. We call such a tree a 𝒴-tree.

𝒴-trees are convenient because they describe the marginal distribution of two leaves in any ultrametric weighted rooted binary tree. Formally:

#### Observation 1

(𝒴-trees provide marginals). Let 𝒯 be an ultrametric weighted rooted binary tree with height *h*. Let *u, v* be two leaves in 𝒯 at distance *t*. Then for any CTMC parameterized by *θ*, the distribution of (*X*_*u*_, *X*_*v*_) under ℙ_*θ, 𝒯, k*_ is the same as the distribution of (*X*_1_, *X*_2_) under ℙ_*θ, 𝒴* (*t,h*),*k*_.

We can now state and prove our general theorem for evolutionary models over ultrametric trees. Since the theorem contains significant amounts of mathematical notation, first we state an informal version of it:

#### Theorem 1 (informal).

If the tree whose topology we want to reconstruct is ultrametric and has known height, and if the parameters of the data generating process are known, then it is possible to reconstruct the tree topology with a small number of characters *k*. The algorithm consists of first computing the function *f*(*t*) which describes the expected (normalized) Hamming distance for two leaves at distance *t*. Next, the empirical raw Hamming distance matrix *D* is ‘corrected’ by using *f*^−1^ to obtain 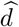, which is an estimate of true tree distance. The root *r* of the tree is treated as a new leaf when constructing 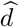. Finally, Neighbor-Joining is applied to 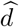, and the resulting unrooted tree is rooted at *r*.

We now state and prove the rigorous version of the theorem:

**Theorem 1** (Probabilistically accurate ultrametric rooted tree reconstruction for parameter-less evolutionary models). Let *h >* 0 be a real number. Let ℳ be an evolutionary model where Θ = {*θ*} is a singleton (i.e. the parameters of the CTMC are known and equal to *θ*), and where 𝕋 is the set of ultrametric weighted rooted binary trees of height exactly *h*. Let *D* be a dissimilarity function, and let *D*_*k*_ be the dissimilarity matrix associated to ℳ_*k*_ and *D*. Suppose that *c* is an upper bound on *D*. Let 𝒜 be an uDBA with *l*_∞_-radius *R*, and let 𝒯 ∈ 𝕋 be any tree (the one whose rooted tree topology we want to recover). Then, for any *δ* ∈ (0, 1], if the number of characters *k* is large enough such that

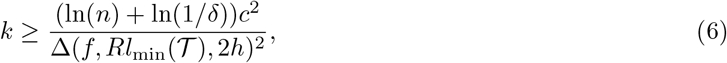

then running 𝒜 on the corrected distances 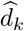– which we define below – and rooting the resulting tree at *r*(𝒯) gives the correct rooted tree topology of 𝒯 with probability at least 1 − *δ*.

The corrected distances *d*_*k*_ are obtained as follows. Define *f* : [0, 2*h*] → ℝ_≥0_ to be the expected dissimilarity for two leaves at distance *t*:

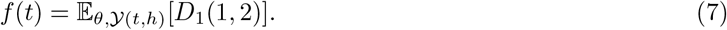

We assume that *f* is continuous, strictly increasing, and *f*(0) = 0. The matrix of corrected distances 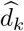 is then the random dissimilarity matrix over *L*(𝒯) ∪ {*r*(𝒯)} defined as:

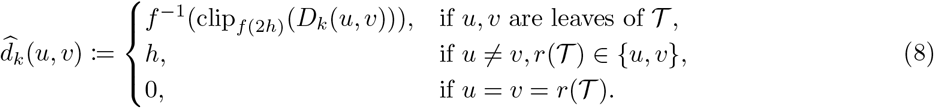

*Remarks*:

1. We assume ultrametric trees of known height because this is the typical setup in CRISPR-Cas9 lineage tracing [13]. However, it is easy to generalize our framework to non-ultrametric trees given additional constraints on the underlying CTMC; we give this result in *SI Appendix, SI Text* B in Theorem 7.
2. Using the shorthand Δ = Δ(*f, Rl*_min_(𝒯), 2*h*), note that Δ scales proportionally to *c*, in the sense that if *D* is multiplied by some factor *κ*, then Δ increases by *κ* too. Thus *c*^2^ and Δ^2^ in the bound (6) cancel each other out, making the bound independent of the units of *D* (as expected).
3. Our bound (6) has the classical form that arises in statistical phylogenetics, with ln(*n*) dependence on the number of leaves *n* and ln(1*/δ*) dependence in the failure probability *δ*; see for example [23, 24, 25, 27, 28, 29, 19].
4. As we will see in our concrete bounds for CRISPR-Cas9, the dependence of Δ on *l*_min_(𝒯) will be of the classical form 1*/l*_min_(𝒯)^2^, again mimicking what is known for other models such as NCF and JC [23, 24, 25, 27, 28, 29, 19].
5. Finally, as we will see in our concrete bounds for CRISPR-Cas9, the dependence of Δ on the mutation rate *λ* will essentially be exponential (e.g. *e*^*λ*^), meaning that the larger the mutation rate, exponentially more characters are needed. For some models such as NCF, this exponential dependency on mutation rate can be avoided with additional constraints on the branch lengths [30]. However, it cannot be removed in general.

As mentioned previously, a different formulation of Theorem 1 which allows for non-ultrametric trees and enables theoretical guarantees for models such as NCF and JC as corollaries is given in *SI Appendix, SI Text* B. This illustrates the generality of our framework.

All that remains is some analytical tool to control the minimum increment for functions *f* that arise in applications. The mean value theorem immediately implies the following simple, concise bound:

#### Lemma 2

(Gradient bound). Let *d*_max_ *>* 0 be a real number. Let *f* : [0, *d*_max_] → ℝ_≥0_ be strictly increasing and differentiable. Then for any 0 *< a* ≤ *d*_max_ and *τ* ∈ [0, *a*], we have the bound:

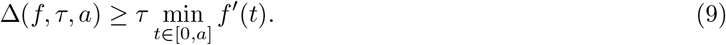

The following lemma establishes an upper bound on the minimum increment of the expected hamming distance *f* for the CRISPR-Cas9 evolutionary model:

#### Lemma 3

(Expected Hamming distance *f* for the CRISPR-Cas9 evolutionary model). For the CRISPR-Cas9 evolutionary model, we have

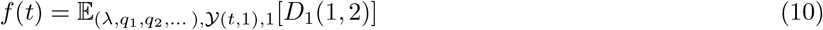

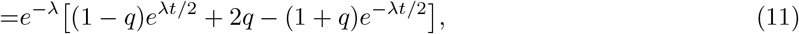

and furthermore, if *q <* 1 and 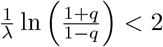, then, for any *τ* ∈ [0, 2],

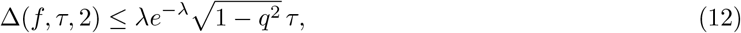

and otherwise

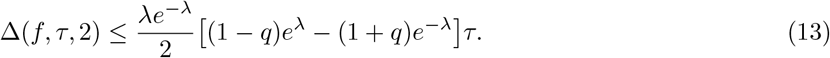

We are now ready to derive concrete bounds on *k* for the CRISPR-Cas9 evolutionary model when using NJ applied to the corrected distances:

#### Theorem 2

(Theoretical guarantees for the CRISPR-Cas9 model with known parameters). Let ℳ_*k*_ be the CRISPR-Cas9 evolutionary model with *k* similarly evolving characters with known parameters, meaning Θ = {*θ*} where *θ* = (*λ, q*_1_, *q*_2_, …) is known. Let *D* be the indicator for equality, so that *D*_*k*_ is the average Hamming distance. Define 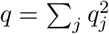 to be the *collision probability*. Let 𝒯 ∈ 𝕋 be any tree (the one whose rooted tree topology we want to recover). Then, for any *δ* ∈ (0, 1], if *q <* 1 and 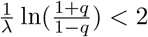, then whenever the number of characters *k* is large enough such that

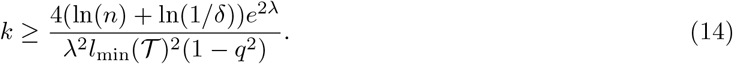

Neighbor Joining run on 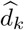– which we define below – and rooting at the root *r*(𝒯) will return the correct rooted tree topology of 𝒯 with probability at least 1−*δ*. Otherwise, in the extreme case *q* = 1 or 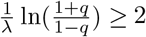, then whenever

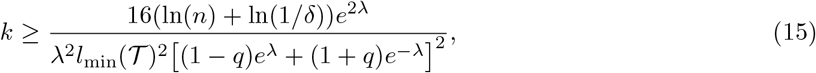

running Neighbor Joining on 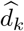 and rooting at the root *r*(𝒯) will return the correct rooted tree topology of 𝒯 with probability at least 1 − *δ*.

The corrected distance matrix 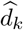 is constructed as follows. Take *f* : [0, 2] → ℝ_≥0_ to be

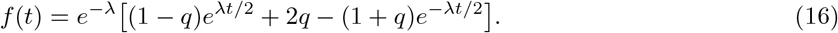

Then *f* is continuous, strictly increasing, and *f*(0) = 0. Define the dissimilarity matrix 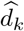 as

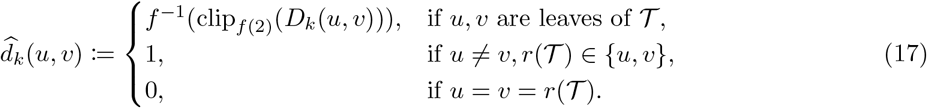

*Remark*. Our bounds are quantitatively similar to those derived recently by [19] for character-based algorithms for the CRISPR-Cas9 model. Importantly, we have (i) a logarithmic dependence on the number of samples *n*, (ii) logarithmic dependence on the inverse recovery error 1*/δ*, (iii) inverse quadratic dependence on the minimum edge length *l*_min_(𝒯), (iv) bounds that tend to +∞ as the mutation rate becomes too low (*λ* → 0) or too high (*λ* → + ∞). For other models such as NCF, such dependencies on tree height and minimum edge length are known to be important and in some cases unavoidable [30].

### 4.2 Analysis of distance-correction scheme with missing data

Many evolutionary models are affected by missing data. For example, sequencing dropouts and transcriptional silencing (heritable missing data) pervade CRISPR-Cas9 lineage tracing data [8, 31, 13, 14]. In protein alignments, gaps are commonly modeled as ignorable missing data [32]. When there is missing data, some of the entries of the character matrix *X*_*L*(𝒯)_ may be hidden from us. We formalize this as in Rubin’s work [33, 34] by augmenting the evolutionary model with a missing data mechanism. To be precise:

#### Definition 10

(Evolutionary model with missing data). Consider the setting of Definition 1. An *evolutionary model with missing data* is defined as follows. In addition to the stochastic process ℙ_*θ,𝒯, k*_ over the tree T thus defined so far, there is a random *missing data mask M* ∈ {0, 1}_*n×k*_ specifying which entries of the character matrix are observed; *M*_*ij*_ = 1 if entry (*i, j*) of the character matrix is observed, and *M*_*ij*_ = 0 otherwise. The joint distribution of the stochastic process over 𝒯 and the missing data mask *M* is specified via the conditional probability distribution of the missing data mask given the stochastic process, denoted by *g*_*ϕ*_(*M*|·). Here *ϕ* denotes the parameters of the missing data mechanism (if any), taking values in some set Φ. The allowed combination of parameters (*θ, 𝒯, ϕ*) is given by a set Γ Θ ⊆ × 𝕋 × Φ. Given *g*_*θ*_ and Γ, the *evolutionary model with missing data* (with *k* similarly evolving characters) is defined as the statistical model

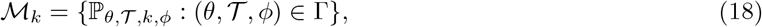

where ℙ_*θ, 𝒯, k,ϕ*_(*M*, ·) := ℙ_*θ, 𝒯, k*_(·)*g*_*ϕ*_(*M*|·). In particular, the joint distribution of *M* and the character matrix is given by ℙ_*θ, 𝒯, k,ϕ*_(*X*_*L*(𝒯)_, *M*) = ℙ_*θ, 𝒯, k*_(*X*_*L*(*𝒯*)_)*g*_*ϕ*_(*M*|*X*_*L*(*𝒯*)_). The *observed* character matrix is no longer *X*_*L*(𝒯)_ but rather constructed by taking *X*_*L*(*T*)_ and replacing each entry where *M*[*i, j*] = 0 with a −1. Formally, the observed character matrix is now 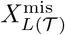 where:

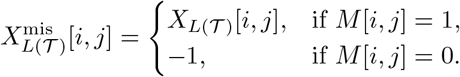

Note that this definition is fully general, and in particular the distribution of *M* may be dependent on *X*. Our theoretical results, however, will hold only when *M* is independent from *X*, as explained later. Allowing Γ to be a strict subset of Θ × 𝕋 × Φ enables the missing data mechanism to depend on the model parameters, such as on the tree 𝒯.

For the CRISPR-Cas9 model, we will consider the following evolutionary model with missing data:

#### Definition 11

(CRISPR-Cas9 evolutionary model with missing data). Consider the setting of Definition 6. We will extend the model to produce missing data as follows: (1) via heritable missing data, wherein at a given rate *r*_silencing_ a character in a cell may become epigenetically silenced during the experiment, and will thus be missing in every descendent of the cell, and (2) via RNA-sequencing dropouts, wherein each entry of the character matrix goes missing i.i.d. with some probability *p*_dropout_. This is a sophisticated missing data mechanism that depends not only on *p*_dropout_ and *r*_silencing_ but also on the tree 𝒯, thus *ϕ* = (*p*_dropout_, *r*_silencing_, 𝒯), Φ = [0, 1] × ℝ_≥0_ × 𝕋 and Γ = {(*θ*, 𝒯, *ϕ*) : *θ* ∈ Θ, 𝒯 ∈ 𝕋, *ϕ* ∈ Φ}.

The combination of the two sources of missing data of the CRISPR-Cas9 model imply that any given entry in the character matrix will be missing marginally with probability 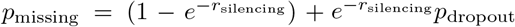. Note that due to heritable missing data, the entries of the missing data mask *M* are not independent. Despite its complexity, the missing data mechanism of the CRISPR-Cas9 model satisfies the important property that *M* and *X*_*L*(𝒯)_ are independent, since we can think about generating *M after* generating *X*_*L*(𝒯)_ and *without* using the value of *X*_*L*(𝒯)_. Indeed, heritable epigenetic silencing and sequencing dropouts can be generated retrospectively, by first generating the lineage tracing data *X* without missingness, and then retrospectively analyzing on which branches of the tree heritable epigenetic silencing would have occurred, as well as which barcodes would have been dropped out during RNA-sequencing. Importantly, this retrospective generation of missing data does not use the values in *X*, which is why *M* and *X* are independent for the CRISPR-Cas9 model with missing data, although *M* and *X*^mis^ are not. One can thus think about *X* as a counterfactual: the lineage tracing data that would have been generated had there been no epigenetic silencing nor sequencing dropouts. This property of independence between *M* and *X* is known as the MACAR (missing always completely at random) condition, and implies that the distribution of the observed entries of the character matrix are the same as their distribution when there is no missing data:

#### Definition 12

(MACAR condition) An evolutionary model with missing data is said to satisfy the MACAR condition if *M* is independent from *X*_*L*(𝒯)_.

Fortunately, it is easy to adapt Theorem 1 to the case when the evolutionary model with missing data satisfies the MACAR condition, as in the CRISPR-Cas9 setting. To derive bounds, we will let *p*_both obs_ *>* 0 be a real number which is a lower bound on the probability that for a fixed pair of leaves (*u, v*), and any character *i*: 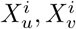 are both *not* missing. For example, in the case of the CRISPR-Cas9 model, our prior lab’s work [19] shows that missing data between two different leaves is positively correlated due to heritable epigenetic silencing and therefore we may take *p*_both obs_ = (1 − *p*_missing_)^2^.

To adapt the algorithm in Theorem 1 to deal with missing data, we extend the definition of *D*_*k*_ as follows:

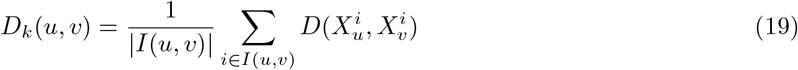

where *I*(*u, v*) ⊆ {1, 2, …, *k*} is the set of characters that are both *not* missing in leaf *u* and *v*. If *I*(*u, v*) is empty, let *D*_*k*_(*u, v*) be 0, so that it is well-defined. With this definition, the algorithm in Theorem 1 is now well-defined for evolutionary models with missing data.

Now, we explain how the proofs and bounds change in Theorem 1, assuming that the MACAR condition is met. The key step that fails in our proofs when there is missing data is the Hoeffding bound in (84). Indeed, *D*_*k*_ is no longer an average of *k* values, but of a *random* number of values, which may thus be much smaller that *k*. As a consequence, using the shorthand Δ = Δ(*f, Rl*_min_(𝒯), 2*h*), the following inequality used in our proofs is *no longer justified* for the chosen value of *k*:

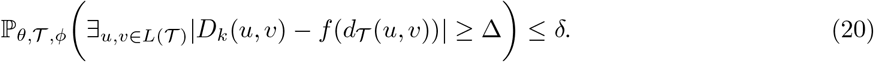

Let *k*_original_(*δ*) be the value of *k* required by the original theorems to achieve an error at most *δ*, namely

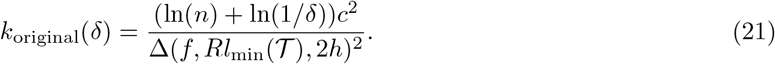

As mentioned, (20) fails for *k* = *k*_original_(*δ*). We will construct *k*_missing_(*δ*), a new value of *k* that ensures that (20) holds when there is missing data. To do this, we choose *k*_missing_(*δ*) such that

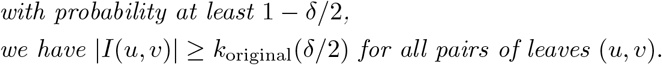

If we can ensure this, we will be done, because with probability at least 1 − *δ/*2 (20) will hold, and this bound holds with probability at least 1 − *δ/*2 for all pairs of leaves, so overall, the algorithm will succeed with probability at least 1 − *δ*. Formally, and using the shorthand Δ = Δ(*f, Rl*_min_(𝒯), 2*h*), we are done because, denoting by *A* the event ∃_*u,v*∈*L*(*𝒯*)_|*D*_*k*_(*u, v*) −*f*(*d*_*𝒯*_ (*u, v*)) ≥ Δ, and *B* the event ∀_*u,v*∈*L*(*𝒯*)_|*I*(*u, v*)| ≥ *k*_original_(*δ/*2) we have:

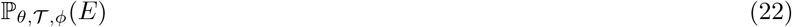

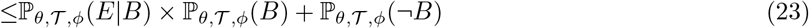

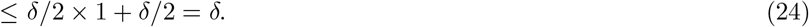

To ensure condition 22, we just need to control a binomial tail bound, for which we use the following result:

#### Lemma 4

(Binomial tail bound). Let *Y* be a binomial random variable with success probability *p*, and suppose that we want *k* successes with probability at least 1 − *ϵ*. Then, it suffices to take the following number of trials 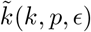:

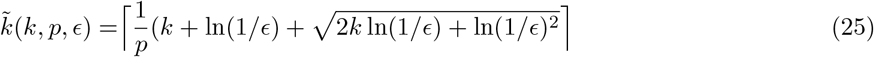

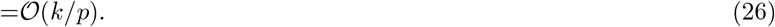

Moreover, if *k* ≥ ln(1*/ϵ*) then we can take the simpler (but weaker in terms of constants):

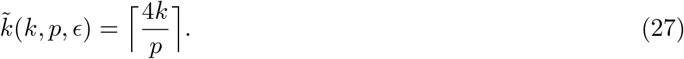

With this, we are ready to state the theoretical reconstruction guarantees for evolutionary models with missing data:

#### Theorem 3

(Bounds for evolutionary models with missing data). In Theorems 1 and 7 (and therefore, in Theorem 2 and Corollaries 3, 4), suppose that the models are equipped with a missing data mechanism as in Definition 10, and suppose that this missing data mechanism is MACAR, meaning that *M* and *X*_*L*(*𝒯*)_ are independent for all (*θ, 𝒯, ϕ*) ∈ Γ. Let *k*_original_(*δ*) be the number of characters to fail with probability at most *δ* assuming there is no missing data. For the model with missing data, suppose that *p*_both obs_ is a lower bound on the probability that 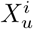 and 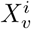 are both observed, and is valid for all *u, v, i*. Then, if *D*_*k*_ is defined as in (19), the theorems hold with the following adjusted value of *k*:

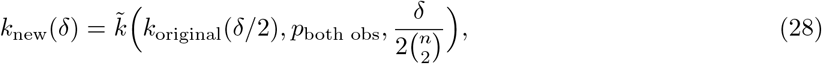

where 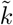 is defined as in (25). Moreover, we can take the simpler (but weaker in terms of constants):

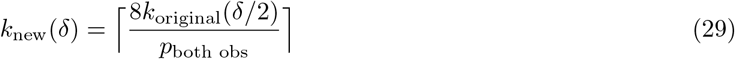

Quantitatively, note that the adjusted *k*_new_(*δ*) is of the order of *k*_original_(*δ/*2)*/p*_both obs_ (when using the stronger bound from (25)), so that one roughly needs 1*/p* times more characters. This make sense, since for example if 50% of the terms in the distance *D*_*k*_ are missing, it means that we need approximately twice as many characters as originally, which recovers the lost terms due to missing data.

In the case of the CRISPR-Cas9 model where each entry of the character matrix is missing i.i.d. with probability *p*_missing_, we get:

#### Corollary 1

(CRISPR-Cas9 with known parameters and missing data). In Theorem 2, let *k*_original_(*δ*) be the number of characters to fail with probability at most *δ*. Then, for the CRISPR-Cas9 model with missing data as in Definition 11, when *D*_*k*_ is defined as in (19), then Theorem 2 holds with the following adjusted value of *k*:

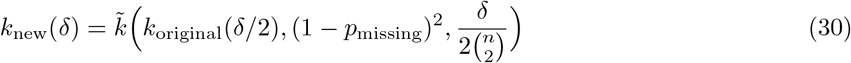

where 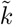 is defined as in (25) and 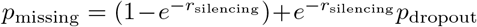. In fact, the simpler expression for 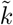 may be used, that is to say, we may take:

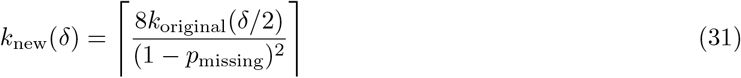

so that the number of characters is 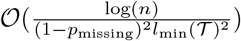.

This settles the case of missing data within our framework. We now move on to the more challenging problem of deriving theoretical guarantees when the model contains unknown parameters, i.e. when *θ* is not known. Importantly, this means that *f* is not known, so that the algorithms described so far cannot be applied.

### 4.3 Analysis of distance-correction scheme with unknown model parameters

Theorem 1 requires knowledge about the parameters of the CTMC. This may be approximately correct for some models such as the JC model, but for the CRISPR-Cas9 evolutionary model - which is our model of interest - this is not the case in practice. Importantly, while the collision probability 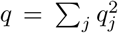 can in principle be estimated by pooling over data from many distinct experiments (since they all share the same *q*) [13, 17], the mutation rate parameter *λ* is unique to each experiment and must be estimated from the character matrix from that experiment alone. Therefore, in this section we tackle the problem of proving theoretical tree reconstruction guarantees using Neighbor-Joining (or any uDBA with a known *l*_∞_-radius) for evolutionary models when the parameters of the model may not be known. This is a novel technical contribution that exceeds the scope of prior theoretical work in statistical phylogenetics, since the theoretical results that are known there only apply to models with known parameters like the NCF or JC model [23, 24, 25, 30, 19]. As before, the minimum increment functional of the true (unknown) function *f* will play a key role in our theory.

We provide a general theorem that can be easily specialized to different evolutionary models, and use it to provide theoretical reconstruction guarantees for the CRISPR-Cas9 evolutionary model when the model parameters are not known. The key obstacle to overcome is that when the parameters of the model are not known, the expected raw dissimilarity function *f*(*t*) is not known, and therefore the algorithm in Theorem 1 (and Theorem 7) cannot be applied. To tackle this, we will assume access to an *approximate version of f in the l*_∞_*-norm*; the algorithm will use this approximate version of *f* instead. The key contribution is deriving theoretical guarantees when this approximate version of *f* is used in the algorithm. As before, these guarantees will depend on the minimum increment functional of the *true* function *f*, but additionally, they will also depend on the *l*_∞_-norm between *f* and the approximation of *f* used. The key lemma we will need to derive these theoretical guarantees is the following, which allows us to control the error introduced by using an approximate version of *f* in the algorithm; this is the counterpart of Lemma 1 where there is error in *f* rather than error in *D*:

#### Lemma 5

Let *d*_max_ *>* 0 be a real number. Let 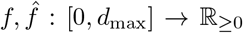 be continuous strictly increasing functions with 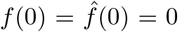. Let *τ* ∈ [0, *d*_max_]. Suppose that 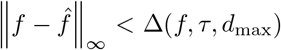. Then, for any *y* ≥ 0 we have that

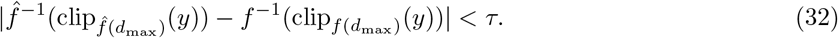

We are now ready to prove theoretical guarantees for evolutionary models with unknown parameters. As before, we consider ultrametric trees as in the CRISPR-Cas9 setting, but more general results are possible; a variation that applies to models such as CFN and JC is provided in Theorem 8. Intuitively, the proof just uses the triangle inequality to to separately control the impact of the error in *f* and the error in *D* towards 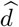. We first state an informal version:

#### Theorem 4 (informal).

In Theorem 1, when the parameters of the data generating process are not known, we can still obtain reconstruction guarantees provided that we are able to estimate the expected dissimilarity function *f* well enough in the *l*_∞_-norm. The algorithm proceeds by using the estimated *f* instead of the unknown, true *f*.

Now formally:

#### Theorem 4

(Probabilistically accurate unrooted tree reconstruction for evolutionary models with unknown parameters over ultrametric trees). Let *h >* 0 be a real number. Let ℳ be an evolutionary model where 𝕋 is the set of ultrametric weighted rooted binary trees of height *h*. Let *D* be a dissimilarity function, and for all *k*, let *D*_*k*_ be the dissimilarity matrix associated to ℳ_*k*_ and *D*. Suppose that *c* is an upper bound on *D*. Let 𝒜 be an uDBA with *l*_∞_-radius *R*. Let *θ* ∈ Θ be the true, *unknown* value of the CTMC’s parameter. Define *f* : [0, 2*h*] → ℝ_≥0_ to be the expected dissimilarity for two leaves at distance *t*:

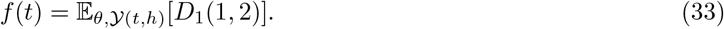

Assume that *f* is continuous, strictly increasing, and *f*(0) = 0. Let 𝒯 ∈ 𝕋 be any tree (the one whose rooted tree topology we wish to recover), and let *δ >* 0 be the tolerated error for accurate reconstruction. Let 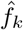 be a continuous, strictly increasing function with 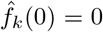 that is an estimate of *f* based on the character matrix *X*_*L*(*𝒯*)_ from ℳ_*k*_. Suppose that there is some 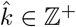 that satisfies:

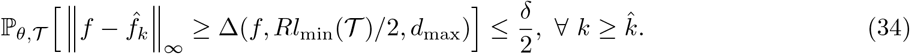

Define the random dissimilarity matrix 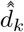 over *L*(𝒯) ∪ {*r*(𝒯)} as:

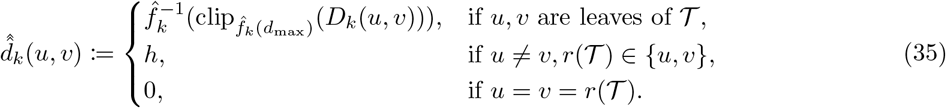

(Note that 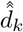 is an estimate of the unknown 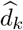 since we do not know *f*, which is why we chose the double-hat notation.) Then, if the number of characters *k* is large enough such that

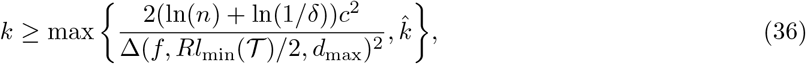

then running 𝒜 on 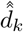 and rooting the resulting tree at *r*(𝒯) gives the correct rooted tree topology with probability at least 1 − *δ*.

A natural way to construct 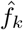 is to use some estimator 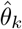 for *θ* with data from ℳ_*k*_, and define:

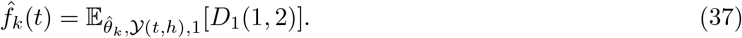

This is the approach we will use for CRISPR-Cas9. Our key result is the following theorem:

#### Theorem 5

(Theoretical guarantees for the CRISPR-Cas9 model with unknown parameters). Let ℳ_*k*_ be the CRISPR-Cas9 evolutionary model with *k* similarly evolving characters with unknown parameters, meaning that Θ = {(*λ, q*_1_, *q*_2_, …) : *λ >* 0, *q*_*j*_ ≥ 0 ∀ *j, Σ* _*j*_ *q*_*j*_ = 1}. Let *D* be the indicator for equality, so that *D*_*k*_ is the average Hamming distance. Define 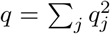 to be the *collision probability*. Let A be an uDBA with *l*_∞_-radius *R*. Let 𝒯 ∈ 𝕋 be any tree (the one whose rooted tree topology we want to recover). Then, for any *δ* ∈ (0, 1], if the number of characters *k* is large enough such that

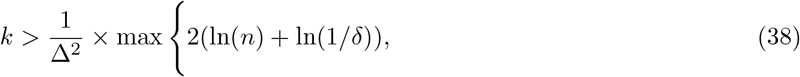

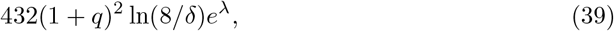

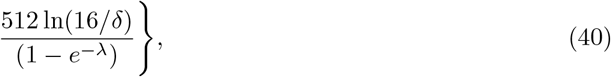

then Neighbor-Joining run on 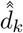– which we define below – and rooting at *r*(𝒯) will return the correct rooted binary tree topology of *T* with probability at least 1 − *δ*. Plugging in *R* = 1*/*2 for Neighbor-Joining and our estimates of Δ from Lemma 3 provides lengthy but concrete bounds, whose asymptotics are still 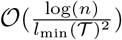 in *n* and *l*_min_(𝒯). Concretely, if *q <* 1 and 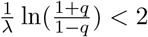, then whenever:

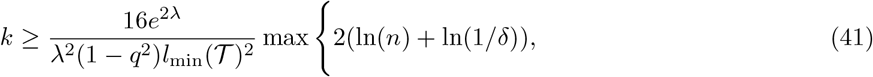

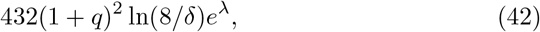

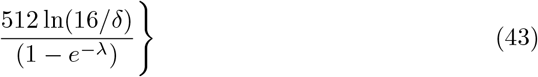

Neighbor Joining run on 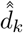 and rooting at the root *r*(𝒯) will return the correct rooted tree topology of 𝒯 with probability at least 1 − *δ*. Otherwise, in the extreme case *q* = 1 or 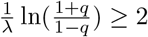, then whenever

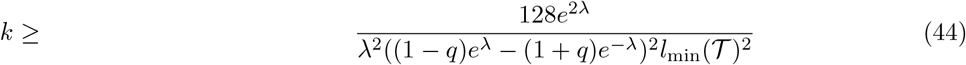

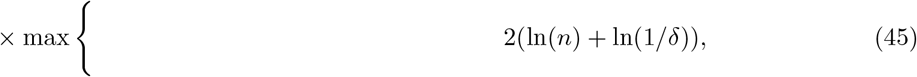

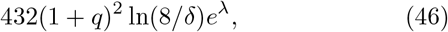

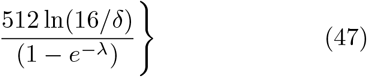

running Neighbor Joining on 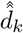 and rooting at the root *r*(𝒯) will return the correct rooted tree topology of 𝒯 with probability at least 1 − *δ*.

We now explain how to construct the corrected distance matrix 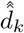. For *λ, q >* 0, let *f* _*λ,q*_: [0, 2] → ℝ_≥0_ be the true expected Hamming distance function:

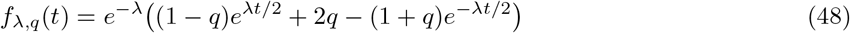

Then *f*_*λ,q*_ is continuous, strictly increasing, and *f*(0) = 0. Let *p*_*k*_ be the following natural estimator of *e*^−*λ*^ (which is the probability of no mutation at a site) based on the *k* i.i.d. observations of the fixed leaf 1:

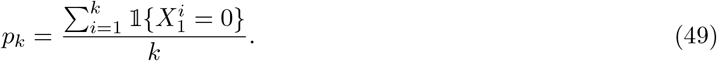

(Note that in practice, one would use all entries of the character matrix, but our result is easier to prove this way.) Let 1 ≤ *M*_1_ *< M*_2_ *<* · · · *< M*_*L*_ ≤ *k* denote the indices of the mutated (i.e. non-zero) characters in leaf 1; note that *L* is a random variable as are the *M*_*i*_. Define the following natural estimator of *q* (which importantly is essentially unbiased since the characters are independent):

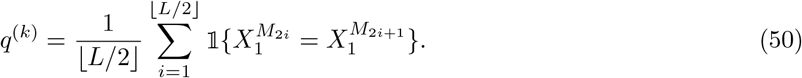

If *L* = 0, define *q*^(*k*)^ = 0 instead. (Again, note that in practice, one would use all entries of the character matrix, but our result is easier to prove this way.) Define the random function 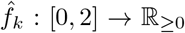: [0, 2] → ℝ_≥0_, which is an estimate of *f*_*λ,q*_, as

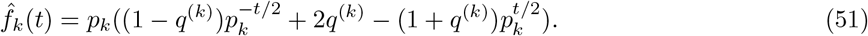

If *p*_*k*_ = 0, arbitrarily set *p*_*k*_ = 1 in (51) so that 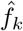 is always strictly increasing and our algorithm below is well-defined. We define the dissimilarity matrix of corrected distances 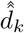 over *L*(𝒯) ∪ {*r*(𝒯)} as:

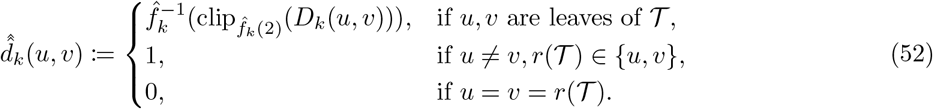

Compared to the bounds with known parameters in Theorem 2, the bounds are similar except for the case when *λ* becomes very large or very small, in which case the second or third term in the maximum will dominate, respectively. The second term in the max accounts for error in estimating *e*^−*λ*^; when *λ* is very large *e*^−*λ*^ is hard to estimate. The third term in the maximum accounts for estimation error in *q*. This error will dominate when *λ* is very small, since 1− *e*^−*λ*^ in the denominator tends to 0. Intuitively, when the mutation rate is so small, there are very few mutations so it is hard to estimate *q*. In this way, we observe that, asymptotically (1) when *λ* is very large, the second term in the max will dominate (i.e. estimation error for *e*^−*λ*^), (2) when *λ* is too small, estimation error of *q* will dominate, and (3) in intermediate regimes the first term (union bound over all leaves) will dominate. Of course, the minimum increment Δ appears in all these terms as 1*/*Δ^2^ and is therefore a key quantity that mediates reconstruction probability.

Finally, note that when there is missing data, the same observation as in Section 4.2 applies to Theorem 5, therefore we obtain:

#### Corollary 2

(CRISPR-Cas9 with unknown parameters and missing data). In Theorem 5, let *k*_original_(*δ*) be the number of characters to fail with probability at most *δ*. Then, for the CRISPR-Cas9 model with missing data as in Definition 11, when *D*_*k*_ is defined as in (19), then Theorem 5 holds with the following adjusted value of *k*:

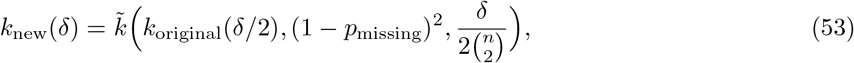

where 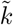 is the function defined in (25) and 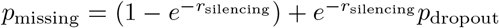. In fact, we may plug in (27), i.e.,

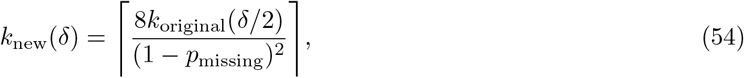

so that the number of required characters is 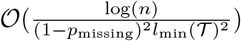.

A concrete, practical algorithm in pseudocode (with the actual estimates *p*_*k*_ and *q*_*k*_ we use in practice) for estimating CRISPR-Cas9 tree topology is provided in *SI Appendix, SI Text* C.

## 5 Experimental Results

### 5.1 Simulated data

We first evaluate the performance of the method on simulated data using the Cassiopeia package [13]. Briefly, our ground truth single-cell phylogenies are simulated under a birth-death process where the birth and death rates are allowed to change with certain probability at each cell division, allowing us to model changes in the fitness of subclades. The simulation ends when a specified population size is reached. With those trees, we then simulate the mutations that are accrued with the Cas9 system. Our simulated lineage tracing assay controls the number of characters, the state distribution *q*_*j*_, the CRISPR/Cas9 mutation rate *λ*, and the rates of missing data. At the end of the simulation, we sample a number of leaves that will be given as input to the algorithm. To explore a wide variety of lineage tracing regimes, we perform the following simulations. We simulate 50 trees with 2000 extant cells each, scaled to have a depth of exactly 1. We then sample exactly 400 leaves from each tree, thus achieving a sampling probability of 20%. The simulation is performed using a birth-death process with rate variation to emulate fitness changes, and is described in detail in *SI Appendix, SI Text* D. The resulting trees display nuanced fitness variation, as can be seen in *SI Appendix*, Fig. 4, which shows 9 of our 50 ground truth induced trees.

Using our simulation framework, we set out to evaluate the performance of our proposed method as judged by the (Camin-Sokal) parsimony score error, Robinson-Foulds, and triplets correct metrics [13]. We further analyze how well the dissimilarity matrix obtained correlates with the ground truth tree distances; we expect the distance correction scheme to improve this correlation.

For each tree, we simulate a lineage tracing experiment with 40 characters. The CRISPR/Cas9 mutation rate is chosen to achieve an expected 50% of mutated entries in the character matrix. We consider 100 possible indel states with a non-uniform probability distribution *q*_*j*_ – as in real data – such that some states are much more common than others. Concretely, the *q*_*j*_ are taken to be the quantiles of an exponential distribution with scale parameter 10^−5^. We introduce sequencing dropouts and epigenetic silencing missing data mechanisms such that on average 20% of the character matrix is missing, with 10% arising from sequencing dropouts and 10% from epigenetic silencing. Epigenetic silencing occurs similarly to CRISPR/Cas9 mutations, happening with a fixed rate during the whole length of the experiment. We call the collection of all these values the ‘default’ lineage tracing parameter regime. For evaluation, we use 50 simulated trees.

We then proceed to repeat the benchmark, this time varying each of the lineage tracing parameters in turn. This allows us to explore lineage tracing datasets with varying levels of quality, as in real data. We vary the number of barcodes in the set {10, 20, 40, 60, 90, 150}, the expected proportion of mutated character matrix entries in the set {10%, 30%, 50%, 70%, 90%}, the number of possible indels in the set {5, 10, 25, 50, 100, 500, 1000}, and the expected missing data fraction in the set {10%, 20%, 30%, 40%, 50%, 60%}, always keeping the expected sequencing missing data fraction at 10%, and adjusting the expected heritable epigenetic missing data fraction accordingly, as we have done in [13].

We benchmark the following four approaches:

1. NJ applied to the Hamming Distance: This is a baseline.
2. NJ applied to the corrected Hamming Distances: This is our proposed improvement over the above baseline, using the corrected distances from Theorem 5.
3. NJ applied to the *weighted* Hamming Distance: This is another baseline. The weighting scheme differs from Hamming Distance in that *D*(*x, y*) = 2 if *x ≠ y* and *x, y >* 0. Therefore, the weighted Hamming distance may take three values: 0, 1 or 2. Weighted Hamming Distance has been observed to perform better on CRISPR-Cas9 data, as compared to standard Hamming distance [16].
4. NJ applied to the corrected weighted Hamming Distances: This is our proposed improvement over the above baseline (NJ applied to the weighted Hamming Distance), using the corrected distances from Theorem 5 as applied to the weighted Hamming distance.

The results for the Robinson-Foulds (RF) metric are shown in Figure 2. The full results are shown in Supplementary Figures 5 and 6. We observe that across essentially all the lineage tracing regimes, using the distance correction scheme provides consistent improvements over the naive version of NJ. Note that in Supplementary Figure 5 lower values of the metric are better, while in Supplementary Figure 6 higher values are better. The best results are obtained using the weighted Hamming Distance with correction (fourth row of the heatmaps). As expected, applying distance correction improves the correlation between the dissimilarity matrix used in NJ and the ground truth tree distance. This may be the underlying cause for the improvement in all other metrics, since the Atteson condition provides reconstruction guarantees for dissimilarity matrices that correlate well with the ground truth distances. Note that although using distance correction provides the best results, there is a big improvement from using a weighted Hamming Distance over a vanilla Hamming distance. For example, remarkably, using the *uncorrected* weighted hamming distance has better Pearson correlation with the ground truth tree distances than the corrected naive Hamming distance. Therefore, distance correction is only as good as the underlying dissimilarity function used. In particular, devising richer dissimilarity functions is a promising direction for furthering the improvements of NJ on CRISPR-Cas9 lineage tracing data.

**Figure 2:**
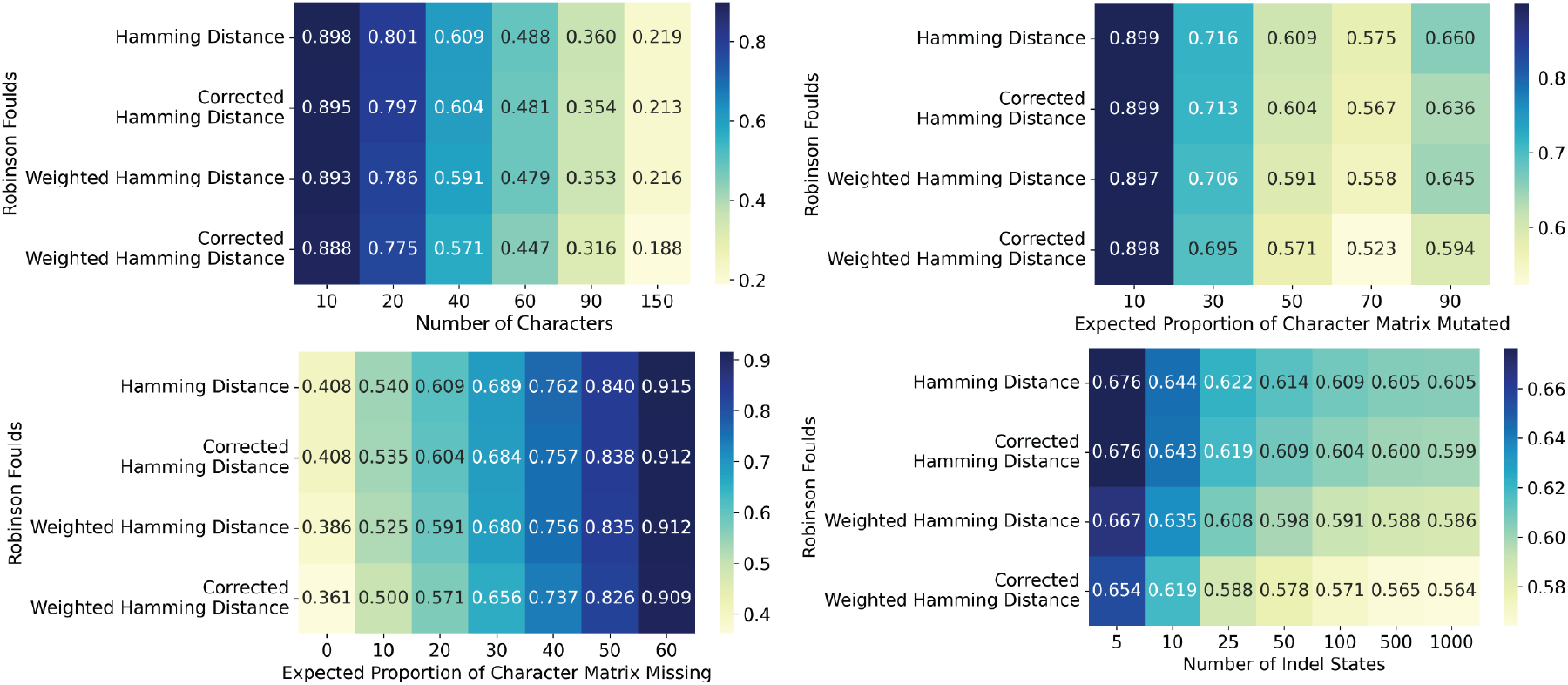
(Using corrected distances improves tree reconstruction on simulated CRISPR-Cas9 data.) We benchmarked two baselines (NJ applied to Hamming Distance, as well as its weighted version) against the corrected versions which we propose. Consistently across the lineage tracing regimes, we see that using distance-correction improves tree reconstruction quality based on Robinson-Foulds. Each entry is the average performance over 50 repetitions.

Finally, we confirmed the consistency of the distance correction scheme by running a simulation where we increased the number of characters until we obtained perfect reconstructions. In this simulation, we considered a fixed perfect binary tree with 512 leaves and equal branch lengths. We used the default lineage tracing regime for simulating CRISPR-Cas9 evolving barcodes, and varied the number of lineage tracing characters in the set of powers of two {4, 8, 16, 32, 64, 128, 256, 512}. The results are shown in Supplementary Figure 7, and are an average over 50 repetitions. All methods achieve perfect performance, which is predicted by our theoretical results in the case of the distance correction scheme.Using raw Hamming distances also achieves consistency, however, for any given fixed number of lineage tracing characters, the distance correction scheme has lower error. In other words, the distance correction scheme has a higher statistical efficiency compared to using raw Hamming distances.

It may be possible to prove the consistency of Neighbor-Joining when using raw Hamming distances, however, our proof techniques will not work. Indeed, our method relies heavily on the Atteson condition, which is not satisfied by the raw Hamming distances, as we have previously discussed. A proof of consistency for Neighbor-Joining using raw Hamming distances may thus need to rely on other results such as quartet-type conditions, but the fact that using raw Hamming distances performs worse than using the distance correction scheme makes it a less attractive route for future work.

### 5.2 Mouse model of lung adenocarcinoma

We next applied our method to lineage tracing data from a mouse model of lung adenocarcinoma [11]. In this work, 21 high-quality clonal populations are analyzed. For each of these, we reconstructed trees using the four aforementioned methods. We excluded the clone *3724 NT T1* since it has over 10,000 cells, which is prohibitive for NJ, leaving 20 clonal populations. We used the value *q* = 0.03 estimated from real data. Duplicate sequences were grouped together prior to tree reconstruction.

Since the ground truth trees are not known for this real dataset, we use the (Camin-Sokal) parsimony score to evaluate the reconstructed trees. The principle is that good reconstructions have better parsimony scores than worse reconstructions, with the ground truth tree being (essentially) the most parsimonious.

The results are shown in Figure 3, where the best method is highlighted for each clone. We can see that on 17 out of the 20 clones (85%), the best results are obtained by using distance correction, and specifically the corrected weighted Hamming distance performs the best. For the other three clones,the vanilla weighted Hamming distance performs best. This shows that the results from simulations translate well to real applications.

**Figure 3:**
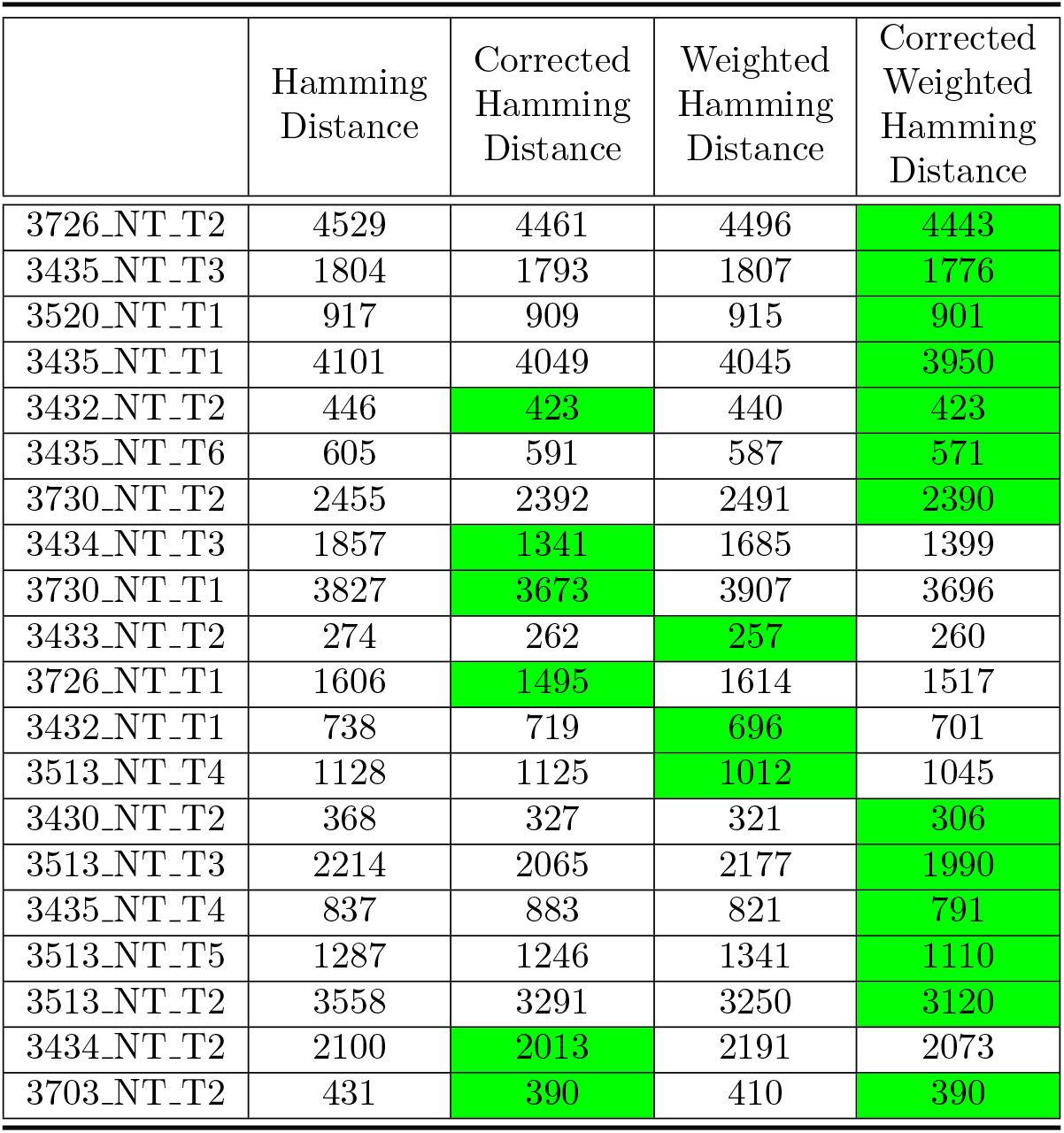
(Distance correction provides the best results on real CRISPR-Cas9 data.) On real data from a mouse model of lung adenocarcinoma [11], our distance correction method improves performance as measured by the (Camin-Sokal) parsimony score of the reconstructed trees. The best method for each clone is highlighted. Specifically, on 17 out of the 20 clonal populations the best results are obtained by using distance correction.

## 6 Discussion

We have developed a general framework to prove theoretical tree reconstruction guarantees for general evolutionary models. Our framework generalizes standard practices in statistical phylogenetics. We applied our framework to the challenging setting of CRISPR-Cas9 lineage tracing data, thereby deriving an algorithm using NJ with theoretical guarantees. Unlike well-studied models such as CFN and JC (for which we also derive theoretical results as a corollary of our general theory), the CRISPR-Cas9 model is complicated by missing data and unknown model parameters. Empirically, we showed that distance-correction scheme improved tree reconstruction quality as measured by diferent metrics, and both on simulated and real data. In general, our framework could be used to derive guarantees for many new evolutionary models in the future, particularly those with missing data and unknown parameters.

It is still important to note that the dissimilarity function used in the scheme is of fundamental importance, to the extent that using a more suitable dissimilarity function may be more important than whether distance correction is used or not. Therefore, devising richer dissimilarity functions for the CRISPR-Cas9 setting is a promising direction of future research that may continue to boost the performance of NJ and more generally of distance-based methods. Note that our distance correction method can be applied to *any* weighted Hamming distance, not just the weightings explored in this work. We expect the best results to be obtained by using such a well-chosen dissimilarity function together with the distance correction scheme proposed in this work. In addition, while we ignore missing data when computing Hamming distances, one may choose to do otherwise. Although this makes it more challenging to compute the expected Hamming distance function and to derive theoretical results – as a detailed model of missing data is now required – it may provide improved empirical results.

Finally, while we consider moment-matching in this work due to its simplicity both in terms of implementation and theoretical analysis, other estimators of pairwise distance such as maximum likelihood estimates (MLE) or regularized versions thereof (as in MAP or posterior mean estimates) may yield further improved results. Indeed, maximum likelihood estimates are statistically efficient provided that the statistical model is well-specified, and may thus outperform moment-matched estimates of pairwise distance. However, theoretical analysis of maximum likelihood estimates in finite samples is challenged by the fact that they involve the maxima of a complicated likelihood function, while moment-matched estimates involve inverses of a simple function (the expected Hamming distance function). The asymptotics of MLE may be easier to derive by bounding the Fisher information, and our intuition is that it would probably provide similar bounds to those we have derived in this work for moment-matching (albeit with better empirical performance). We leave these avenues for future work.

## Supplementary Information

### SI Text

#### A Proofs of the results in the main text

##### A.1 Proof of Lemma 1

Suppose |*y*^*′*^ − *f*(*t*)| *<* Δ(*f, τ, d*_max_). First note that:

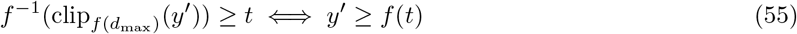

Indeed, since *f* : [0, *d*_max_] → ℝ_≥0_ is a continuous, strictly increasing function with *f*(0) = 0, then 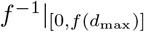 is defined and is strictly increasing, hence:

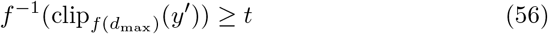

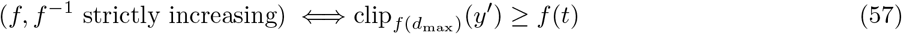

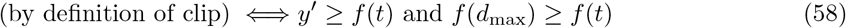

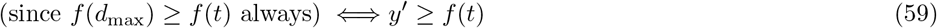

Now we do some casework. If *y*^*′*^ = *f*(*t*) there is nothing to do because 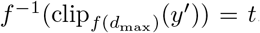. Now, suppose *y*^*′*^ *> f*(*t*). If *t* ∈ [0, *d*_max_ − *τ*] then we have by assumption:

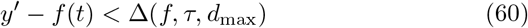

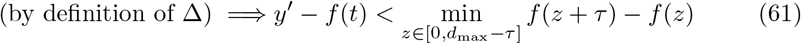

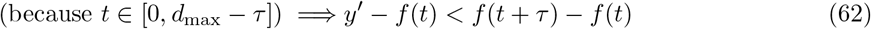

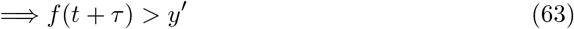

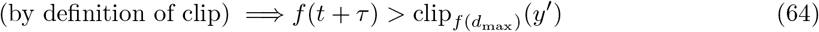

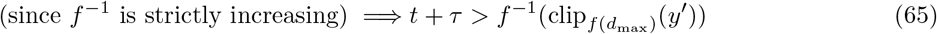

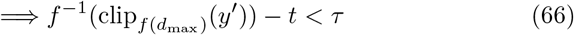

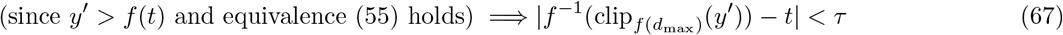

Otherwise, if *t* ∈ (*d*_max_ −*τ, d*_max_] we have 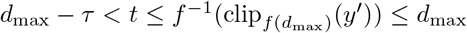 and the claim follows immediately.

Next suppose that *f*(*t*) *> y*^*′*^. If *t* ∈ [*τ, d*_max_] then we have by assumption:

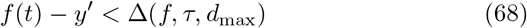

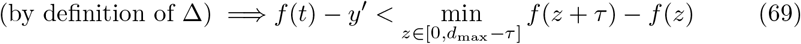

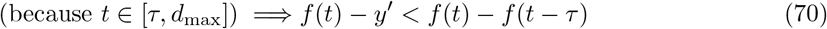

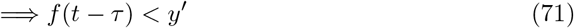

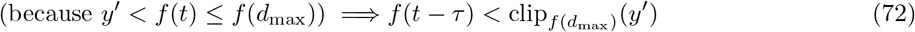

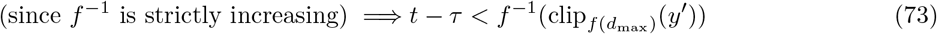

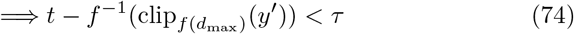

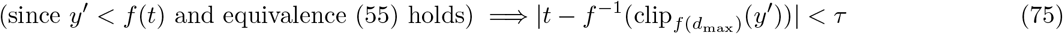

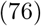

If instead *t* ∈ [0, *τ*) then 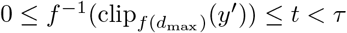 and the claim follows. This completes all casework.

Now we prove the stronger bound. Assume that |*y*^*′*^ −*f*(*t*)| *<* Δ(*f, τ*, min(*d*_max_, *t*+*τ*)). Note that if *t*+*τ > d*_max_ we are done because the statement is equivalent to the original one (since Δ(*f, τ*, min(*d*_max_, *t* + *τ*)) = Δ(*f, τ, d*_max_)). Otherwise, *t* + *τ* ≤ *d*_max_ so that *f*(*t* + *τ*) is well defined and *f*(*t* + *τ*) ≥ *f*(*t*) + Δ(*f, τ, t* + *τ*) by definition of the minimum increment. Since by assumption *y*^*′*^−*f*(*t*) *<* Δ(*f, τ*, min(*d*_max_, *t*+*τ*)) = Δ(*f, τ, t*+*τ*), then combining the two bounds we have: *y*^*′*^ *< f*(*t*) + Δ(*f, τ, t* + *τ*) ≤ *f*(*t* + *τ*) ⇒ 0 ≤ *y*^*′*^ *< f*(*t* + *τ*). Importantly, *y*^*′*^ needs no clipping, i.e. 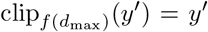. In particular, we can apply *f*^−1^ on both sides to get *f*^−1^(*y*^*′*^) *< t* + *τ*. This yields the sought upper bound on *f*^−1^(*y*^*′*^). For the lower bound on *f*^−1^(*y*^*′*^), suppose by means of contradiction that *f*^−1^(*y*^*′*^) ≤ *t* −*τ*. Then by definition of the minimum increment applying *f* on both sides we have *y*^*′*^≤ *f*(*t*) − Δ(*f, τ, t* + *τ*), contradicting the initial assumption. This way, the stronger bound is proven.

##### A.2 Proof of Proposition 1

Since by assumption A has an *l*_∞_-radius of *R* then by the Atteson condition it suffices to show that for all pairs *u, v* ∈ *L*(𝒯) ∪ {*r*(𝒯)}, we have 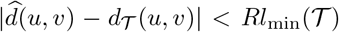. If *u* and *v* are both leaves, then this is equivalent to |*f*^−1^(clip_*f* (2*h*)_(*D*(*u, v*))) − *d*_*T*_ (*u, v*)| *< Rl*_min_(𝒯), which follows immediately by applying Lemma 1 with *t* = *d*_*T*_ (*u, v*), *y* = *f*(*d*_*T*_ (*u, v*)), *y*^*′*^ = *D*(*u, v*), and *τ* = *Rl*_min_(𝒯) (just note that *t* ≤ 2*h* since 2*h* is an upper bound on the diameter of 𝒯). If on the other hand one of *u, v* is equal to the root *r*(𝒯) then 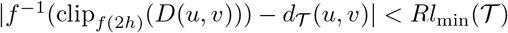 by definition of 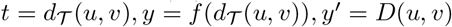, and so 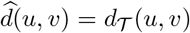 also holds trivially.

##### A.3 Proof of Observation 1

This is true because the CTMC runs down the tree independently over each edge, so it does not matter what the chain is doing on another edge at the same point in time. Thus, the distribution of (*X*_*u*_, *X*_*v*_) depends only on the 𝒴-tree induced by *u, v*, which is equivalent to 𝒴 (*t, h*).

##### A.4 Proof of Theorem 1

It suffices to show that for all pairs of leaves (*u, v*) we have that:

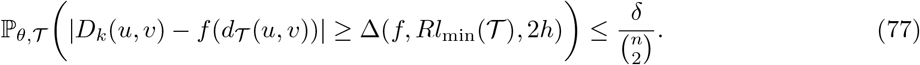

Indeed, if this is the case, then the union bound tells us that with probability at least 1 − *δ* we have, for *all* pairs of leaves (*u, v*): |*D*_*k*_(*u, v*) − *f*(*d*_*T*_ (*u, v*)) |*<* Δ(*f, Rl*_min_(𝒯), 2*h*). Thus, by Proposition 1 we would be done. To show Eq. (77), first note that by Observation 1:

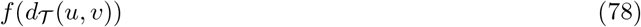

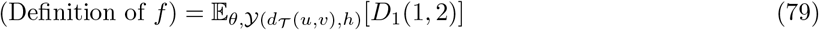

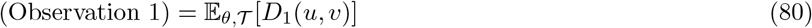

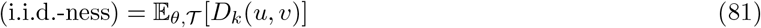

Therefore, applying Hoeffding’s inequality, for a fixed pair of leaves (*u, v*) we have that:

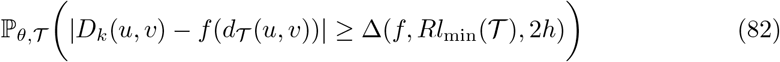

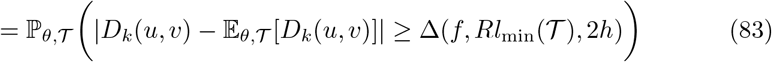

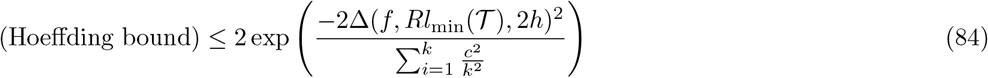

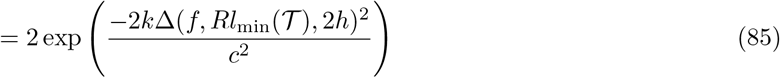

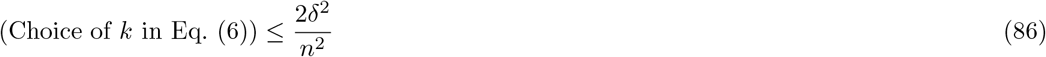

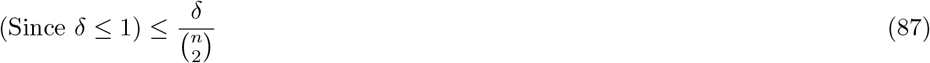

as desired; note that the second to the last inequality is exactly equivalent to Eq. (6).

##### A.5 Proof of Lemma 3

For computing the expectation, note that if *d*_*𝒯*_ (*u, v*) = *t* we have two cases that give us nonzero Hamming distance at a site: either we have that both *u* and *v* each acquired a mutation independently with no collision, or one cell has mutated at that site and the other has not, which can happen in two ways. Both of these need to be multiplied by the probability that there is no mutation on the shared branch. We thus have:

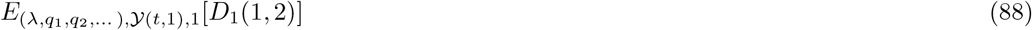

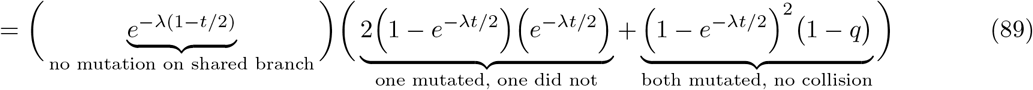

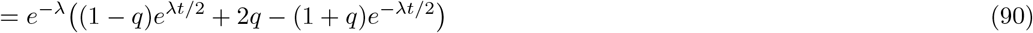

Next, computing some derivatives of *f* we have

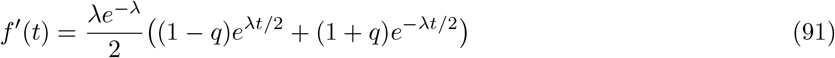

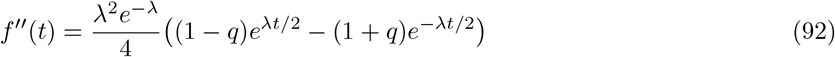

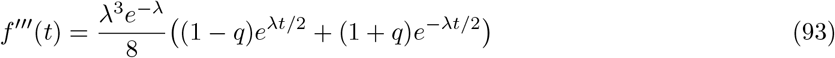

To minimize the first derivative, note that *f*^*′′′*^ *>* 0 and hence *f*^*′*^ is strictly concave, so we set the second derivative to 0 and solving for *t* we get that when *q <* 1, *f*^*′*^ achieves its minimum on ℝ_≥0_ at

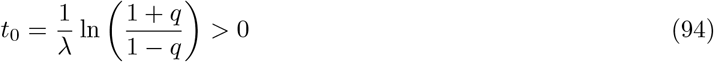

If *t*_0_ ∈ [0, 2], plugging it back in to *f*^*′*^ we get that

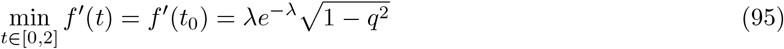

and Eq. (12) follows from the gradient bound of Lemma 2. Otherwise, if *t*_0_ *>* 2 then *f*^*′*^ is minimized at 2 so plugging in *f*^*′*^(2) and applying the gradient bound of Lemma 2 we get Eq. (13). Finally, if *q* = 1 then 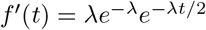 which is also minimized at *t* = 2.

##### A.6 Proof of Theorem 2

Since NJ has an *l*_∞_-radius of 1*/*2 [23], we are done by plugging in *f* and our bound on the minimum increment of *f* from Lemma 3 into Theorem 1.

##### A.7 Proof of Lemma 4

By assumption the number of successes follows a binomial distribution with success probability *p*. For any 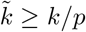, the lower tail Chernoff bound gives us:

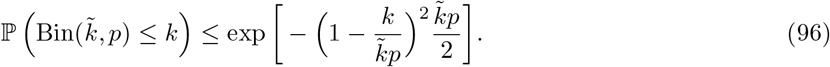

Choosing 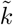 large enough such that the above is at most *ϵ* and solving for 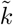 yields Eq. (25). If *k* ≥ ln(1*/ϵ*) Eq. (27) follows immediately from replacing ln(1*/ϵ*) by *k* and upper bounding.

##### A.8 Proof of Theorem 3

As argued above, we only need to prove that with probability at least 1−*δ/*2, for each pair of leaves (*u, v*), the number of characters that are *not* both missing is at least *k*_original_(*δ/*2). For a given pair of leaves, Lemma 4 implies that this condition will be satisfied with probability at least 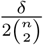. By taking a union bound over all pairs of leaves, we get a failure probability of at most *δ/*2, so we get Eq. (28). To see that the simpler Eq. (29) also works, we just need to use Eq. (27) for which we need to show that 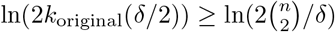; this is trivial using the definition of *k*_original_ and noting that Δ ≤ *c* by assumption that *D* takes values in [0, *c*].

##### A.9 Proof of Lemma 5

First we prove a version of the lemma when there is no clipping:

###### Lemma 6

Let *d*_max_ *>* 0 be a real number. Let *f*, 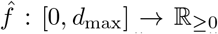be continuous strictly increasing functions with 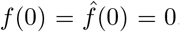. Let *τ* ∈ [0, *d*_max_]. Suppose that 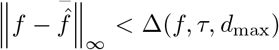. Then, for any *y* ≥ 0such that 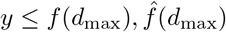 we have that:

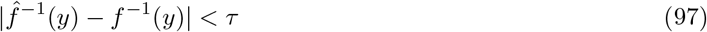

*Proof*. Just apply Lemma 1 with 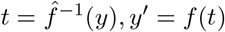 to the function *f*, which tells us that:

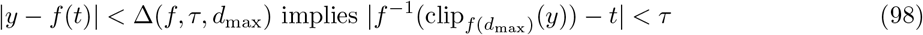

For the term |*y* − *f*(*t*)|, note that:

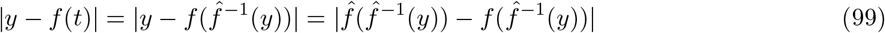

Now,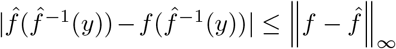, and since by assumption 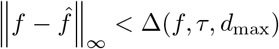 then we sat-isfy the precondition |*y* −*f*(*t*)| *<* Δ(*f, τ, d*_max_) of Eq. (98) and therefore we obtain that 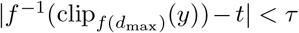. Since by assumption *y* does not clip (i.e. 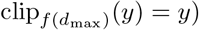, this is equivalent to:

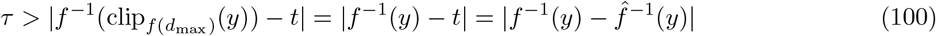

which is exactly what we wanted to show.

Now we prove Lemma 5. If 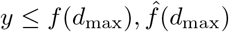 then we are done by Lemma 6. Therefore, we only need to analyze the case when *y* clips for *f* or 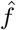, or both. When both of them clip (i.e. 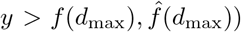, there is nothing to do because then 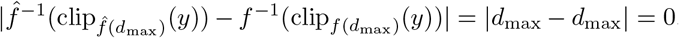 analyze the two remaining cases.

First let’s assume that *f* clips but 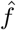 does not, i.e. 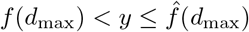. In this case, we have that:

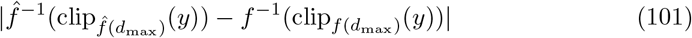

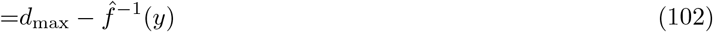

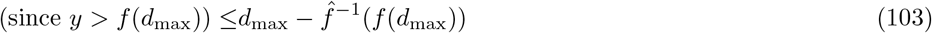

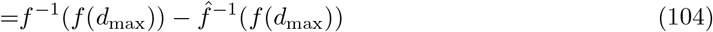

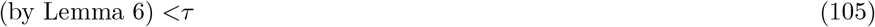

Finally, let’s assume that 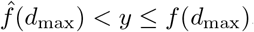. This case is analogous:

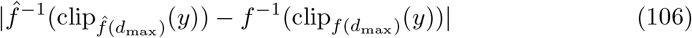

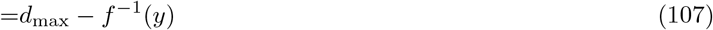

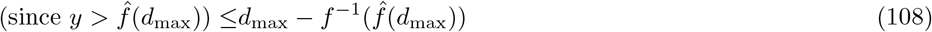

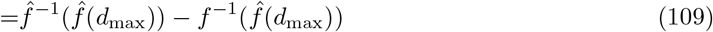

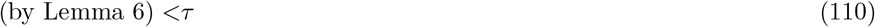

and we are done.

##### A.10 Proof of Theorem 4

Let 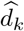 be the dissimilarity matrix when using *f* instead of 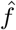 in Eq. (35) (i.e. Eq. (8)). By assumption 34, and since the value of *k* in Eq. (36) is larger than 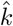, with probability at least 1 − *δ/*2 we have 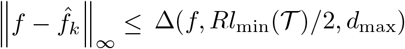, i.e. 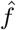 is close to *f*. By Lemma 5 and the definitions of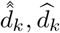, it follows that 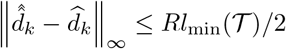 with probability at least 1 − *δ/*2, i.e. 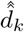 is close to 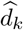. On the other hand, using the same union bound from Theorem 1 (and bounding above to get a clean expression), the value of *k* in Eq. (36) ensures that, with probability at least 1 − *δ/*2, we have

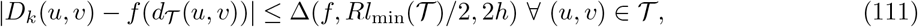

and therefore by applying Lemma 1:

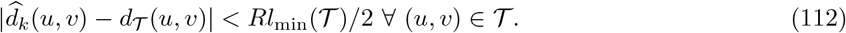

Equivalently, 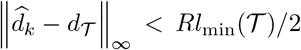. Thus, with probability at least 1 − *δ* we have simultaneously 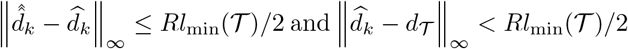, and the triangle inequality implies 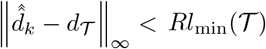.Since 𝒜 has *l*_∞_-radius equal to *R*, we are done.

##### A.11 Proof of Theorem 5

We start by considering the case when only *λ* is unknown, and then give a theorem for the case when both *λ* and the *q*_*j*_ are unknown.

###### Theorem 6

(Theoretical guarantees for the CRISPR-Cas9 model with unknown parameter *λ*). Let ℳ_*k*_ be the CRISPR-Cas9 evolutionary model with *k* similarly evolving characters with unknown mutation rate, meaning that Θ = {(*λ, q*_1_, *q*_2_, …) : *λ* ∈ ℝ_+_} where the *q*_*j*_ are known. Let *D* be the indicator for equality, so that *D*_*k*_ is the average Hamming distance. Define 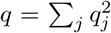 to be the *collision probability*. For *λ >* 0, let *f*_*λ*_ : [0, 2] → ℝ_≥0_ be:

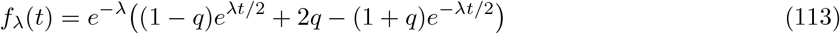

Then *f*_*λ*_ is continuous, strictly increasing, and *f*(0) = 0. Let *p*_*k*_ be the following natural estimator of *e*^−*λ*^ based on the *k* i.i.d. observations of one fixed leaf *u*:

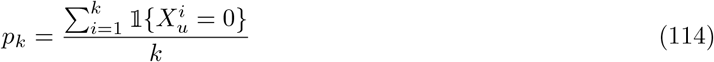

(Note that in practice, one would use all entries of the character matrix, but our result is easier to prove this way.) Define the random function 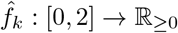, as

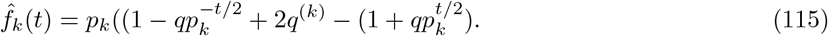

If *p*_*k*_ = 0, arbitrarily set *p*_*k*_ = 1 in Eq. (51) so that 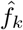 is always strictly increasing and our algorithm below is well-defined. Define the dissimilarity matrix 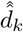 as

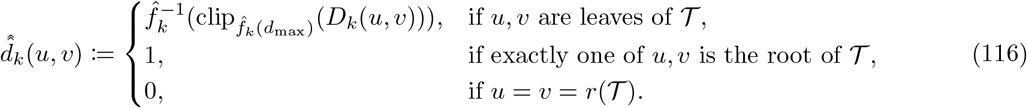

If the number of characters *k* is large enough such that

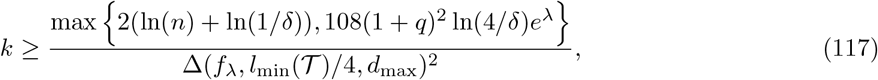

then Neighbor-Joining run on 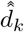 and rooting at *r*(𝒯) will return the correct rooted binary tree topology of *T* with probability at least 1 − *δ*. By using our prior bounds in Lemma 3 for the minimum increment of *f*_*λ*_, we get, more concretely, that if *q <* 1 and 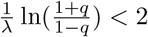, then whenever

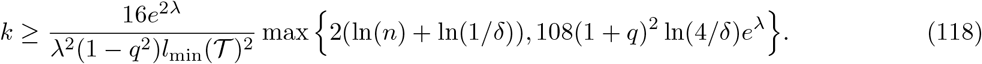

Neighbor Joining run on 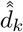 and rooting at the root *r*(𝒯) will return the correct rooted tree topology of 𝒯 with probability at least 1 − *δ*. Otherwise, if *q* = 1 or 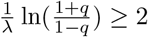, then whenever

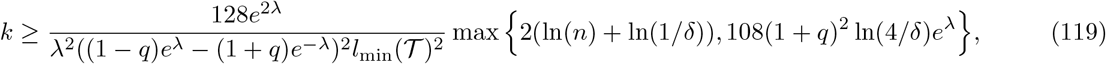

running Neighbor Joining on 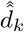 and rooting at the root *r*(𝒯) will return the correct rooted tree topology of 𝒯 with probability at least 1 − *δ*.

*Proof*. In what follows, recall that *R* = 1*/*2. We have already computed *f* and bounded its minimum increment in Lemma 3. Thus, to deal with the unknown parameter *λ*, we just need to find 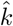 that will achieve:

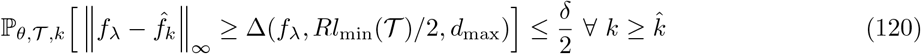

To this end, using the triangle inequality, we have that

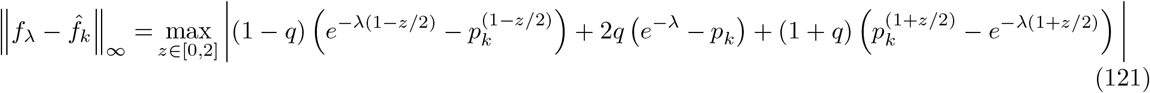

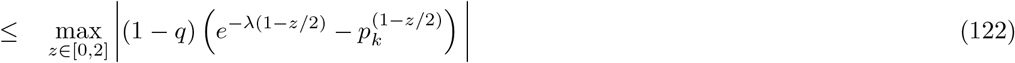

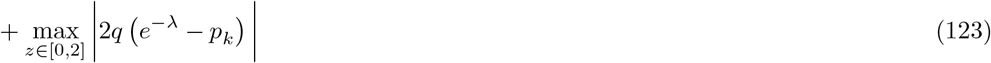

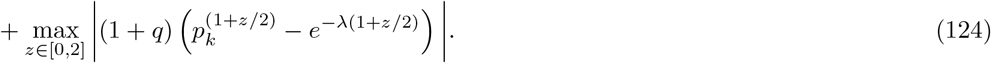

We now use the following lemma to deal with the three summands:

###### Lemma 7

For 0 *< θ* ≤ 1 and 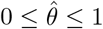 and 0 ≤ *x* ≤ *x*_max_, we have that:

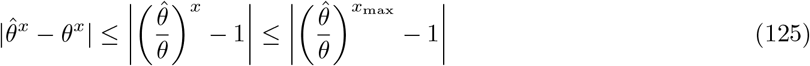

*Proof*. We have

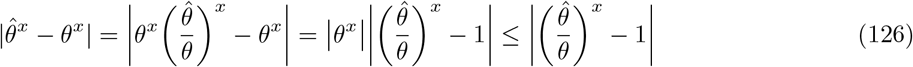

from which the results follows.

Our goal now is to take Lemma 7 with *θ* = *e*^−*λ*^ and 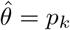 to get an upper bound of Eq. (121) that can later be controlled with a Chernoff bound. We have that:

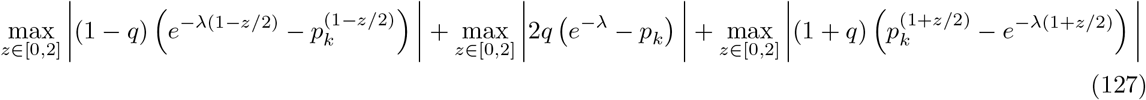

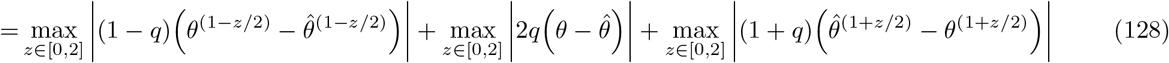

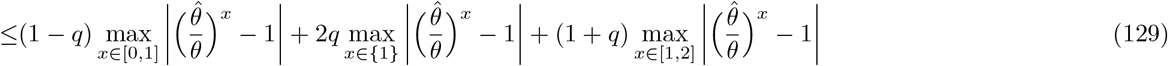

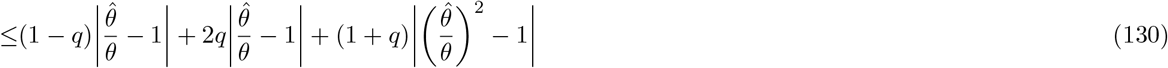

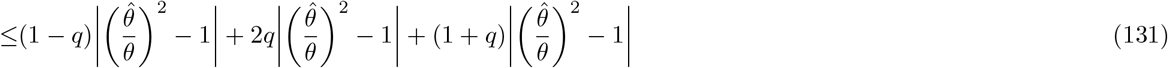

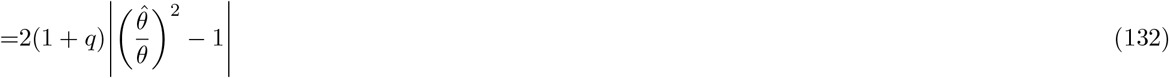

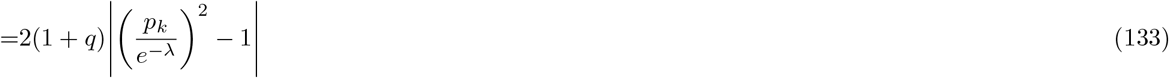

Therefore, to ensure 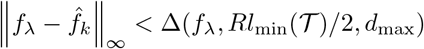 it suffices to have:

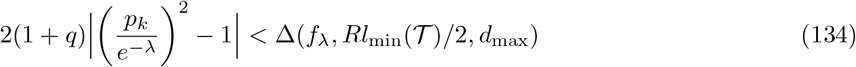

Using the shorthand Δ = Δ(*f*_*λ*_, *Rl*_min_(𝒯)*/*2, *d*_max_), we thus want, with probability at least 1 − *δ/*2:

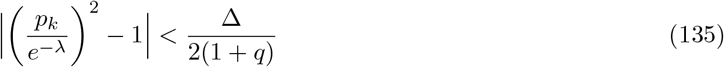

For this, we claim it is in turn sufficient to have, with probability at least 1 − *δ/*2:

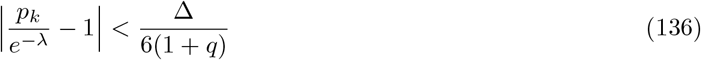

Indeed, if Eq. (136) is true, then:

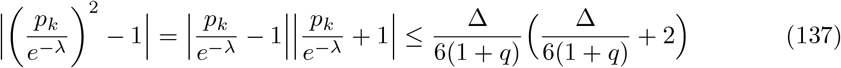

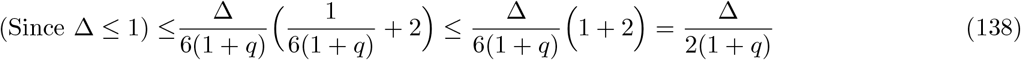

To ensure Eq. (136), taking a multiplicative Chernoff bound of the form:

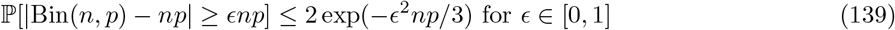

instantiated at 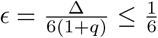 we get that:

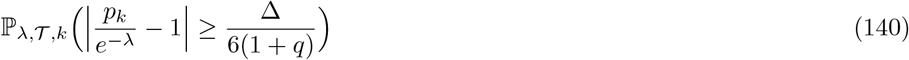

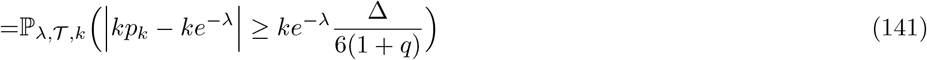

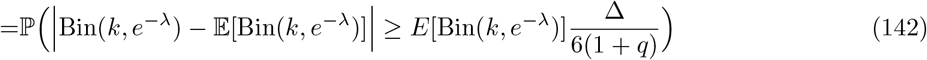

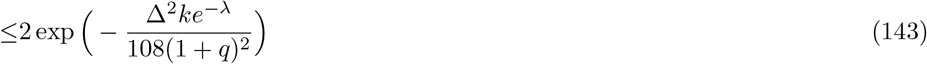

Hence it suffices to take 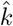 such that:

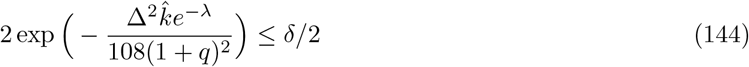

i.e.:

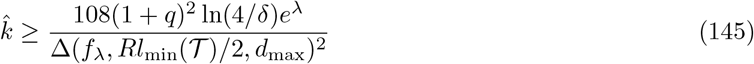

and we are done by Theorem 4.

Now we turn to the case when the *q*_*j*_ are also not known. Note that the algorithm is identical to that in Theorem 4 except that we use the following estimate of *q*: Let 1 ≤ *M*_1_ *< M*_2_ *<*· · · *< M*_*L*_ ≤ *k* denote the indices of the mutated (i.e. non-zero) characters in leaf 1, then:

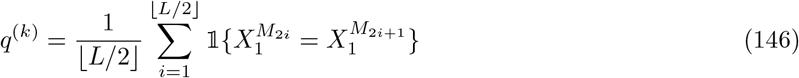

If *L* = 0, define *q*^(*k*)^ = 0 instead. Note that given *L, L >* 0 we have *q*^(*k*)^ ∼ Bin(⌊*L/*2⌋, *q*)*/*⌊*L/*2⌋. Using our machinery, it is a matter of bounding 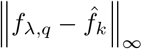 where 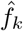 is now defined as:

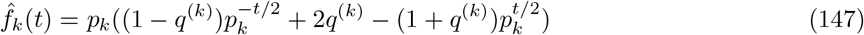

The triangle inequality together with *e*^−*x*^ ≤ 1 for *x* ≥ 0 shows that, compared to our original bound on 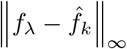, we now have an extra term 4|*q*^(*k*)^ − *q*|, i.e.:

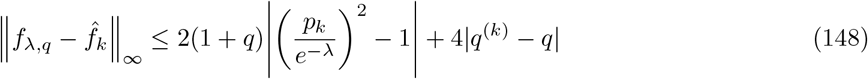

Consequently, using the shorthand Δ = Δ(*f*_*λ,q*_, *Rl*_min_(𝒯)*/*2, *d*_max_), to ensure that:

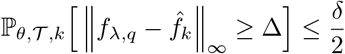

it suffices to have:

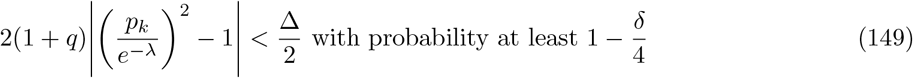

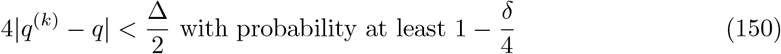

We already know how to ensure Eq. (149) from the proof of Theorem 6, namely with:

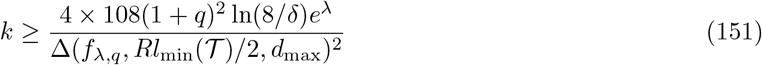

For Eq. (150), the Hoeffding bound yields:

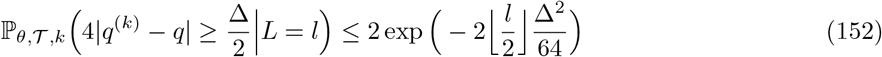

If *L* was fixed and equal to *l*, we would be done with:

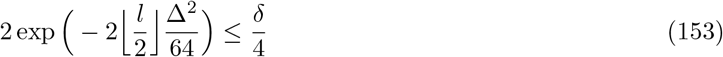

However, *L* is random, with *L*∼ Bin(*k*, 1 − *e*^−*λ*^) so we need to use to use the same technique as for missing data adjustment. Specifically, we split the allowed *δ/*4 error probability into *δ/*8 + *δ/*8, and seek *l* such that we have the more restrictive:

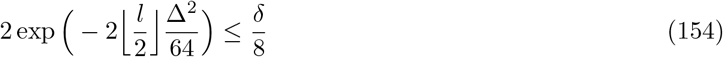

which yields:

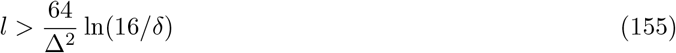

and finally adjust this value of *l* to get that Eq. (150) holds whenever:

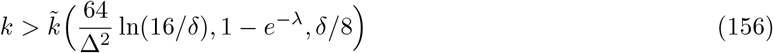

And thus, 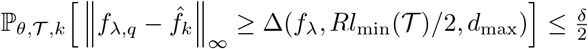 holds for all 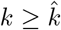 with:

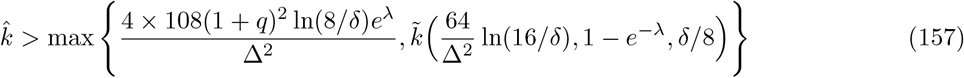

Since 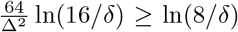 we can use the simpler expression for 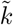, namely 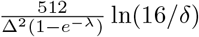. Plugging this value of 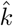 (Eq. (157)) into Eq. (36) in Theorem 4 we are done.

## B Generalization for the CFN and JC models

We first need the analogous proposition to Proposition 1 but for unrooted trees:

### Proposition 2

(Accurate unrooted tree reconstruction assuming pairwise distances can be reasonably estimated through *f*). Suppose that 𝒯 is a weighted unrooted binary tree with diameter at most *d*_max_. Suppose that *D* is a dissimilarity matrix over the leaves of 𝒯, and *f* : [0, *d*_max_] → ℝ_≥0_ is a continuous strictly increasing function with *f*(0) = 0. If 𝒜 is a uDBA with *l*_∞_-radius *R*, and if for every pair (*u, v*) of leaves we have that:

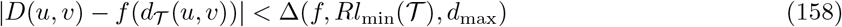

then, letting 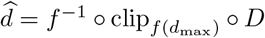, we have that running 𝒜 on 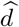 will return the correct unrooted binary tree topology over the leaves of 𝒯 with probability at least 1 − *δ*. Moreover, using the stronger version of Lemma 1, note that Proposition 2 is true with the improved bound:

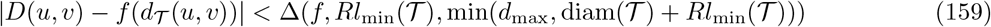

in place of Eq. (158).

*Proof*. Since by assumption 𝒜 has an *l*_∞_-radius of *R* it suffices to show that for all pairs of leaves (*u, v*), we have 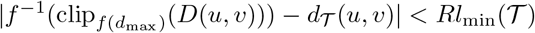. But this follows immediately by applying Lemma 1 with *t* = *d*_*𝒯*_ (*u, v*), *y* = *f*(*d*_*𝒯*_ (*u, v*)), *y*^*′*^ = *D*(*u, v*), and *τ* = *Rl*_min_(𝒯) (just note that *t* ≤ *d*_max_ by the assumption that *d*_max_ is an upper bound on the diameter of 𝒯). For the improved bound, just observe that *t* = *d*_*T*_ (*u, v*) ≤ diam(𝒯) and *τ* = *Rl*_min_(𝒯).

With this, we can prove the following version of Theorem 1 but for stationary, reversible models where rooting does not matter; we highlight the differences with Theorem 1 in blue:

### Theorem 7

(Probabilistically accurate unrooted tree reconstruction for parameter-less stationary reversible evolutionary models over trees of bounded height). Let *h >* 0 be a real number. Let ℳ be an evolutionary model where Θ = {*θ*} is a singleton (i.e. the parameters of the CTMC are known), the CTMC is stationary and reversible, and where 𝕋 is the set of weighted rooted binary trees of height at most *h*. Let *D* be a dissimilarity function, and let *D*_*k*_ be the dissimilarity matrix associated to ℳ_*k*_ and *D*. Suppose that *c* is an upper bound on *D*. Let A be an uDBA with *l*_∞_-radius *R*. Define *f* : [0, 2*h*] → ℝ_≥0_ to be:

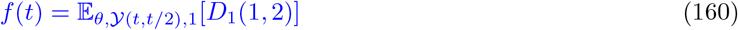

Assume that *f* is continuous, strictly increasing, and *f*(0) = 0. Let 𝒯 ∈ 𝕋 be any tree. Then, if we define the random dissimilarity matrix 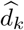 over *L*(𝒯) as:

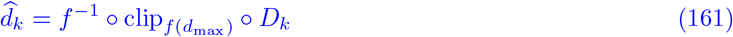

if the number of characters *k* is large enough such that

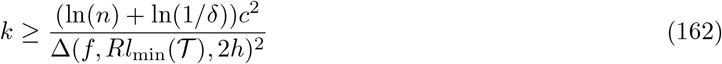

then running 𝒜 on 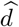 gives the correct unrooted tree topology induced by the leaves of 𝒯, with probability at least 1 −*δ*.

**Remark:** Using the stronger version of Proposition 2, Theorem 7 holds with the improved bound:

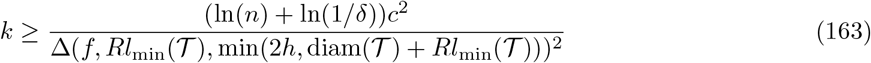

*Proof*. The proof is essentially identical to the proof of Theorem 1. Just note that by stationarity and reversibility of the CTMC:

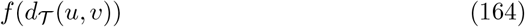

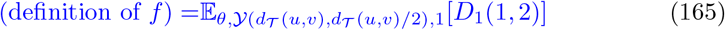

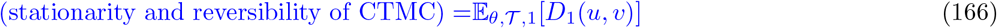

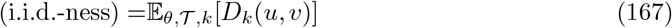

Therefore, applying Hoeffding’s inequality, for a fixed pair of leaves (*u, v*) we have that:

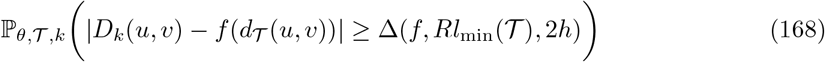

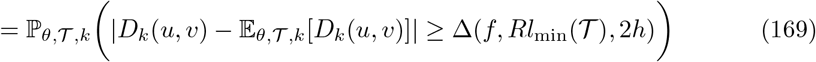

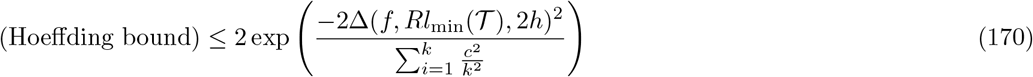

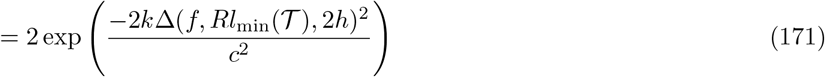

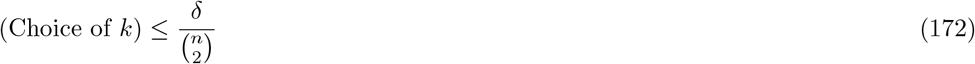

so we are done by the union bound and Proposition 2. The proof of the stronger result uses the stronger version of Proposition 2 instead.

With this version of our framework, we can easily give reconstruction guarantees for NJ applied to the corrected distances under the NCF and Jukes-Cantor models, as is commonly done in statistical phylogenetics and for which similar theoretical guarantees are already known [23, 24, 25, 27, 28, 29]. Our result is as follows:

### Corollary 3

(Theoretical guarantees for the NCF model over trees of bounded height). Let *h >* 0 be a real number. Consider the NCF CTMC, where the starting state is sampled from the uniform distribution over {0, 1} and mutations happen at a rate of 1.0; note that NCF has no parameters, so that Θ = {*θ}* is a singleton (*θ* = 1.0). Suppose that 𝕋 is the set of weighted rooted binary trees of height at most *h*. Let *ℳ*_*k*_ be the evolutionary model with *k* similarly evolving characters defined this way. Let *D*_*k*_ be the average hamming distance matrix. Define

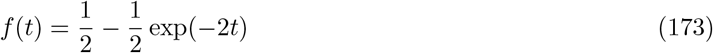

and:

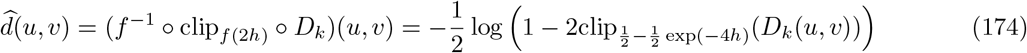

If:

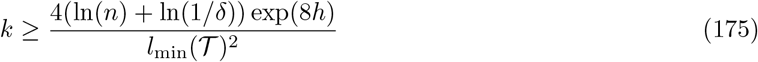

then running Neighbor-Joining on *d* will return the correct unrooted tree topology with probability at least 1 − *δ*. In fact, we have the improved bound which does not depend on *h*:

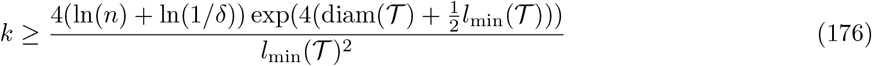

*Proof*. Just apply Theorem 7. As in our guarantees for the CRISPR-Cas9 model, we only need to compute *f* and bound its minimum increment. It is well known that for the NCF model, the probability of being in a different state after time *t* is 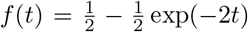. Finally, the minimum increment functional can be bounded via the gradient bound as Δ(*f, τ, a*) ≥ *τ* exp(−2*a*). The stronger tree-dependent bound follows from applying the strengthened version of Theorem 7 instead.

*Remark*. Note that our bound is essentially identical to the known bound of [23] (with *ϵ* in place of 1− *e*^−*ϵ*^ resulting from using the gradient bound as opposed to a manual bound). Thus, our method generalizes the techniques and bounds used in statistical phylogenetics, and we will later apply them to the CRISPR-Cas9 setting, with missing data and unknown parameters.

*Remark*. The stronger bound which does not depend on *h* is helpful because it allows us to set *h* arbitrarily large in the algorithm (basically *h* = +∞), so that the algorithm does not need to have access to a good bound on tree height, all while retaining good tree-specific bounds. Theorem 7 is the reason we are interested in bounds that are tree-dependent.

### Corollary 4

(Theoretical guarantees for the Jukes-Cantor model over trees of bounded height). Let *h >* 0 be a real number. Consider the Jukes-Cantor CTMC, where the starting state is sampled from the uniform distribution over {*A, C, G, T* } and mutations happen at a rate of 1.0; note that JC has no parameters, so that Θ = {*θ*} is a singleton. Suppose that 𝕋 is the set of weighted rooted binary trees of height at most *h*. Let ℳ_*k*_ be the evolutionary model with *k* similarly evolving characters defined this way. Let *D*_*k*_ be the average hamming distance matrix. Define

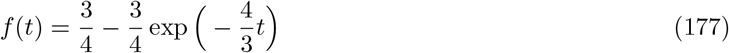

And

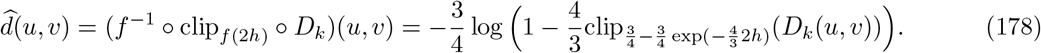

If

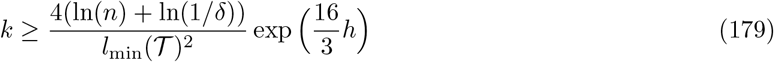

then running Neighbor-Joining on 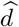 will return the correct unrooted tree topology with probability at least 1 − *δ*. In fact, we have the improved bound which does not depend on *h*:

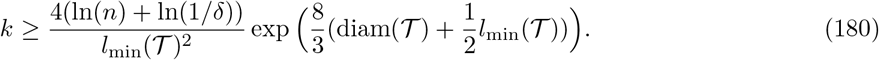

*Proof*. Just apply Theorem 7. As in our guarantees for the CRISPR-Cas9 model, we only need to compute *f* and bound its minimum increment. It is well known that for the Jukes-Cantor model, the probability of being in a different state after time *t* is 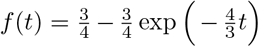. Finally, the minimum increment functional can be bounded via the gradient bound as 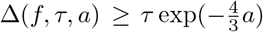. The stronger tree-dependent bound follows from applying the strengthened version of Theorem 7 instead.

Lastly, dealing with unknown model parameters can be done analogously to Theorem 4 (differences in the theorem in blue):

### Theorem 8

(Probabilistically accurate unrooted tree reconstruction for stationary reversible evolutionary models with unknown parameters over trees of bounded height). Let *h >* 0 be a real number. Let ℳ be an evolutionary model where 𝕋 is the set of weighted rooted binary trees of height at most *h*. Let *D* be a dissimilarity function, and for all *k*, let *D*_*k*_ be the dissimilarity matrix associated to ℳ_*k*_ and *D*. Suppose that *c* is an upper bound on *D*. Let A be an uDBA with *l*_∞_-radius *R*. Let *θ* ∈ Θ be the true, *unknown* value of the CTMC’s parameter. Define *f* : [0, 2*h*] → ℝ_≥0_ as follows:

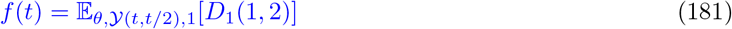

Assume that *f* is continuous, strictly increasing, and *f*(0) = 0. Let 𝕋 be any tree (the one whose rooted tree topology we wish to recover), and let *δ >* 0 be the tolerated error for accurate reconstruction. For all *k*, let 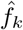 be a continuous, strictly increasing function with 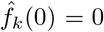 that is an estimate of *f* based on the character matrix *X*_*L*(𝒯)_ from ℳ_*k*_. Suppose that 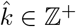 satisfies:

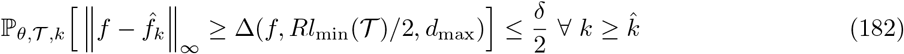

Then, if we define the random dissimilarity matrix 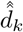 over *L*(𝒯) as:

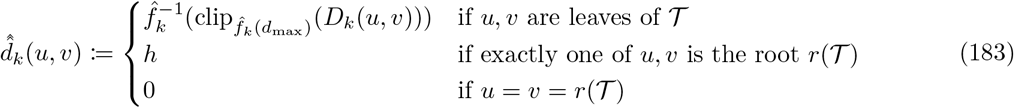

if the number of characters *k* is large enough such that:

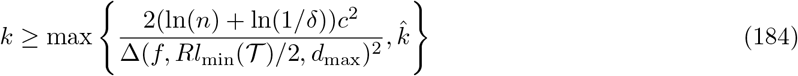

then running 𝒜 on 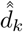 gives the correct unrooted tree topology induced by the leaves of T with probability at least 1 − *δ*.

*Proof*. Analogous to the proof of Theorem 4.

Theorem 8 could be used to derive theoretical guarantees for the Jukes-Cantor model when the transition rates are not all the same and not known. We leave this to future work.

## C Algorithm for CRISPR-Cas9 tree topology estimator

### Algorithm 1

CRISPRCas9TreeTopologyEstimator

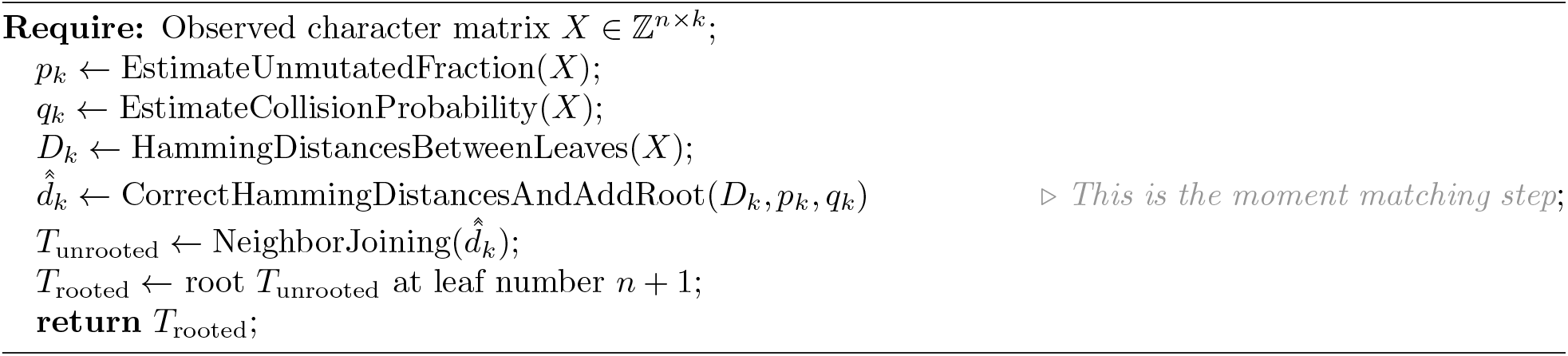

### Algorithm 2

EstimateUnmutatedFraction

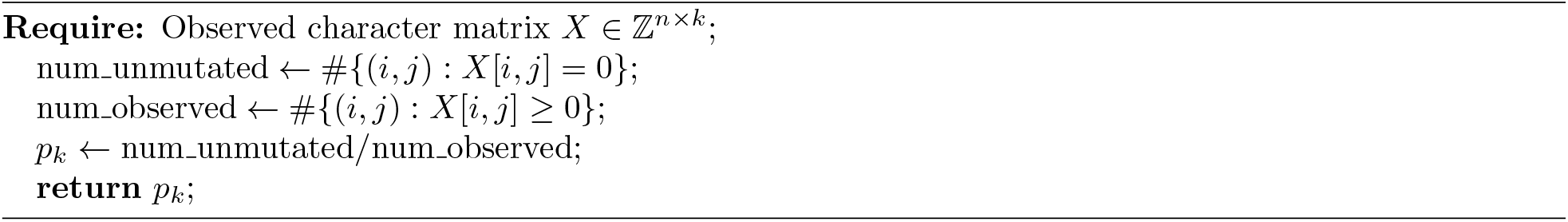

### Algorithm 3

EstimateCollisionProbability

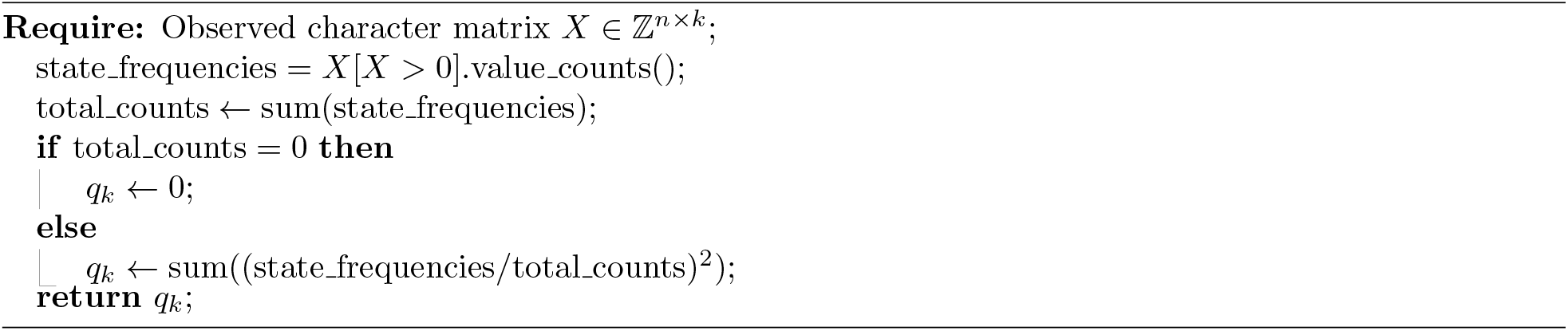

### Algorithm 4

HammingDistancesBetweenLeaves

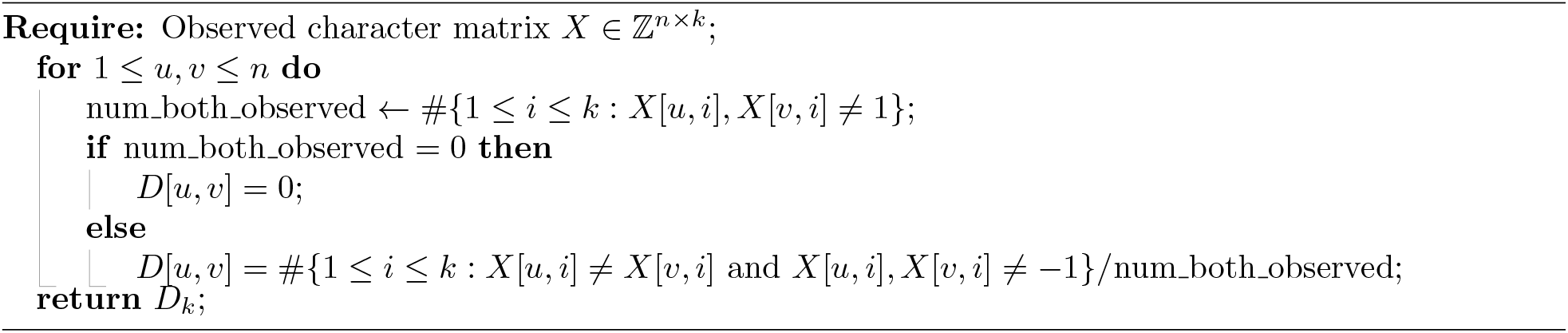

### Algorithm 5

CorrectHammingDistancesAndAddRoot

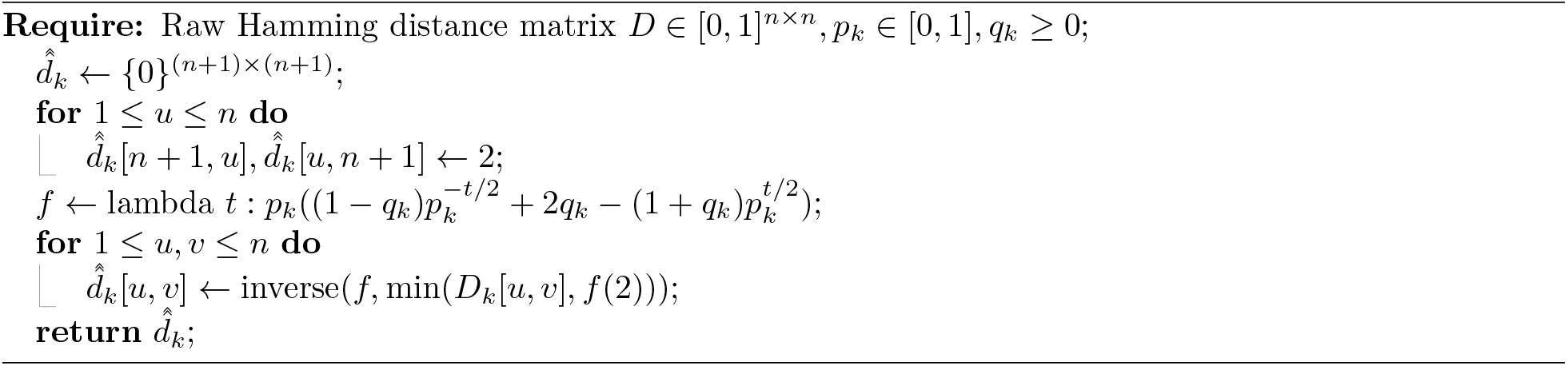

## D Tree Simulation details

Trees were simulated using a subsampled birth-death process. In this simulation, each cell has a birth rate and a death rate. The amount of time before a birth or a death event occurs follows an exponential distribution with the given birth and death rates. Additionally, a cell cannot divide before at least 0.02 units of time have passed, called the *offset*. Initial birth and death rates are set such that birth rate is ten times higher than death rate and such that under a birth-death process with these rates and offset, the expected population size after 1 unit of time is 2000. This yields a birth rate of 11.46 and a death rate of 1.146. Whenever a cell divides, its fitness changes with probability 50%. When such a change in fitness occurs the birth rate becomes is multiplied by 5^*x*^ where *x* is drawn from a Normal distribution with mean 0 and standard deviation 0.5. Once the cell population reaches 2000, we terminate the simulation and sample 400 leaves uniformly at random, to match a sampling probability of 20%. The phylogeny induced by these 400 cells is then the ground truth tree used in the simulations. These trees showcase interesting fitness variation, as displayed in Figure 4.

**Figure 4:**
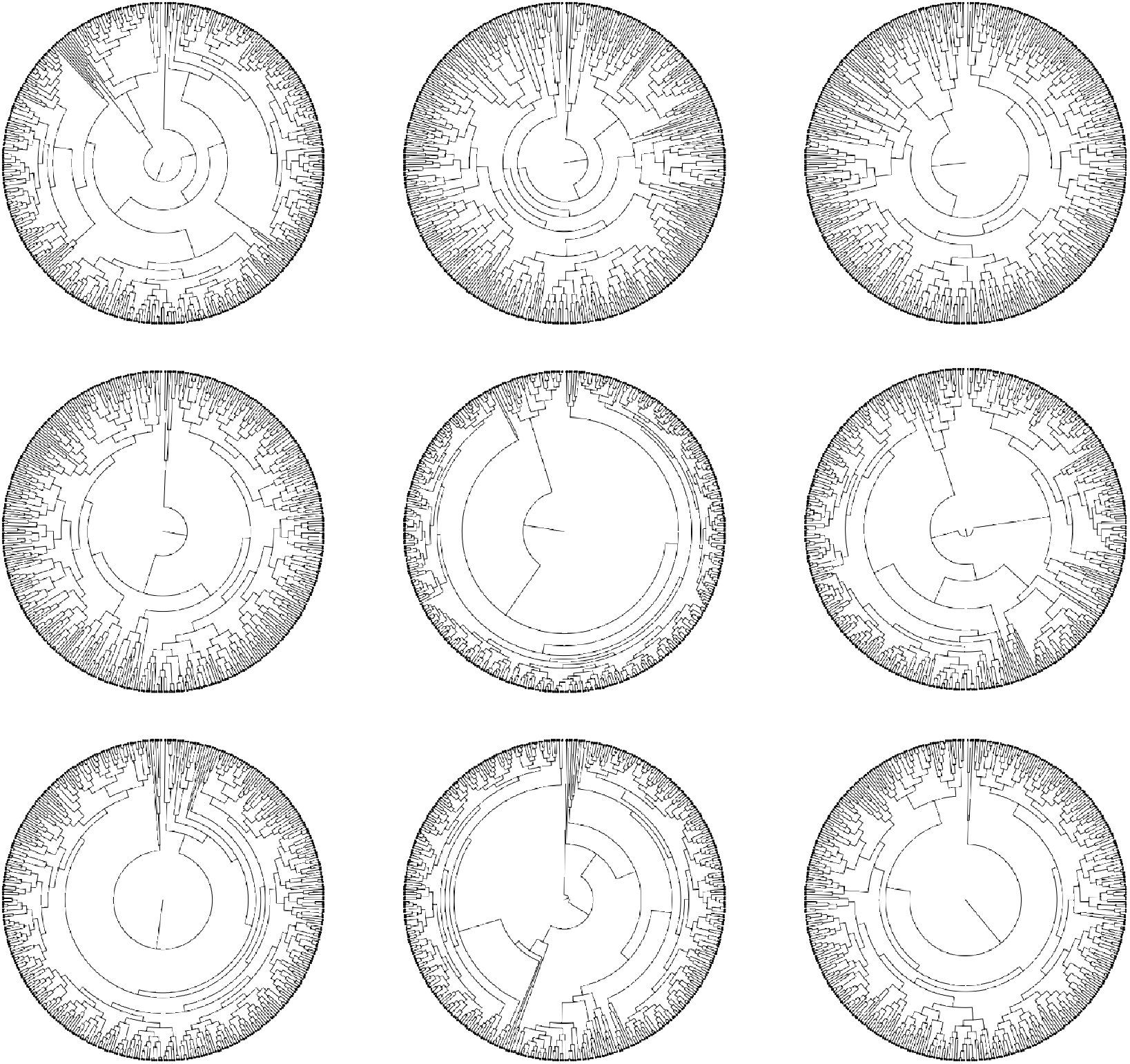
Sample ground truth trees. Ground truth trees corresponding to the first 9 random seeds. Our simulated trees are diverse and showcase subclones with different proliferative capacity.

## Supplementary Figures

**Figure 5:**
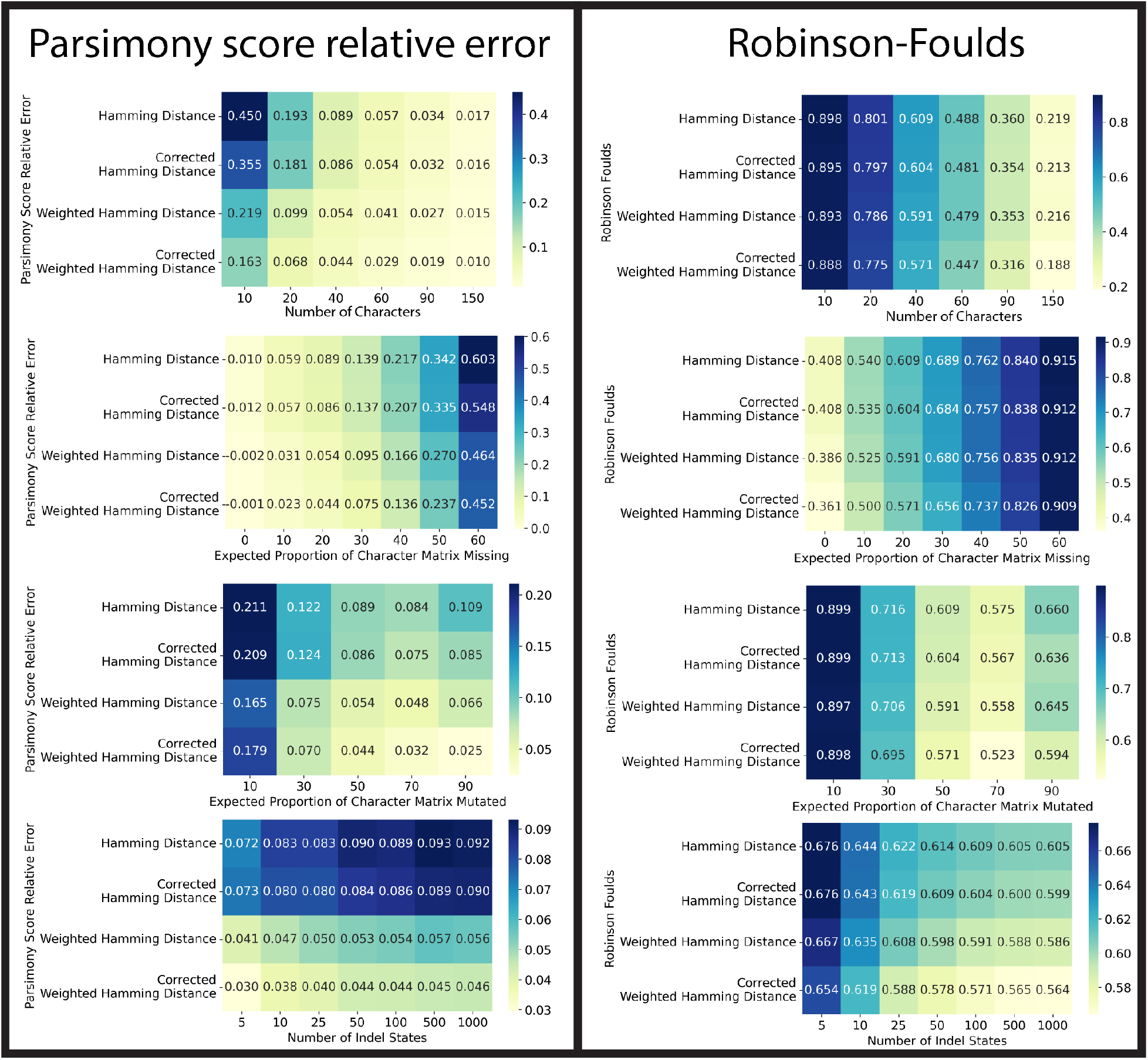
(Using corrected distances improves tree reconstruction on simulated CRISPR-Cas9 data.) We benchmarked two baselines (NJ applied to Hamming Distance, as well as its weighted version) against the corrected versions which we propose. Consistently across the lineage tracing regimes, we see that using distance-correction improves tree reconstruction quality based on (Camin-Sokal) parsimony score and Robinson-Foulds. Each entry is the average performance over 50 repetitions.

**Figure 6:**
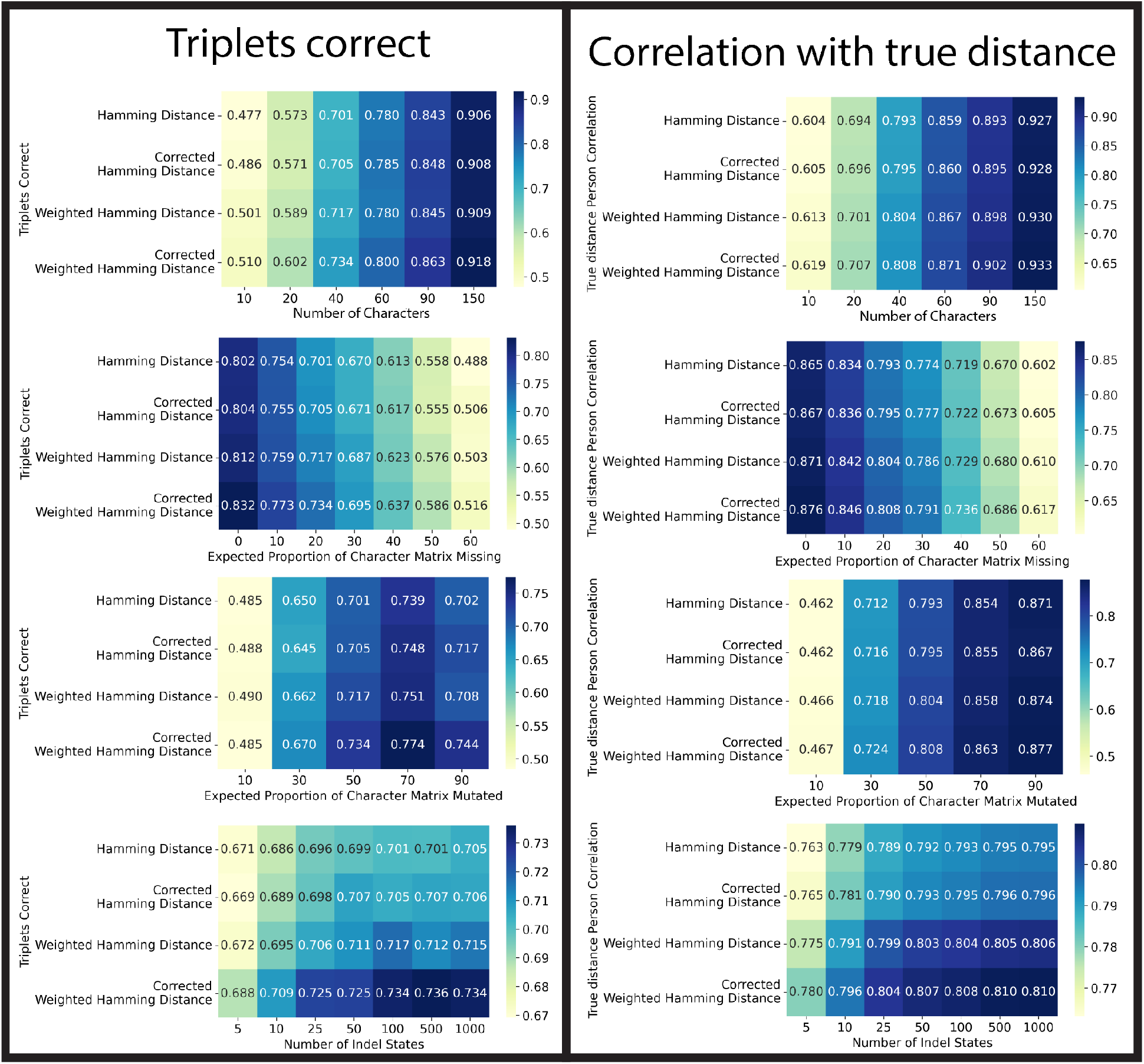
(Using corrected distances improves tree reconstruction on simulated CRISPR-Cas9 data.) We benchmarked two baselines (NJ applied to Hamming Distance, as well as its weighted version) against the corrected versions which we propose. Consistently across the lineage tracing regimes, we see that using distance-correction improves tree reconstruction quality on triplets correct. We hypothesize that the improved Pearson correlation between the dissimilarity matrix used in NJ and the ground truth tree distances (right panel) underlies the improved performance for all other metrics, due to the Atteson condition. Each entry is the average performance over 50 repetitions.

**Figure 7:**
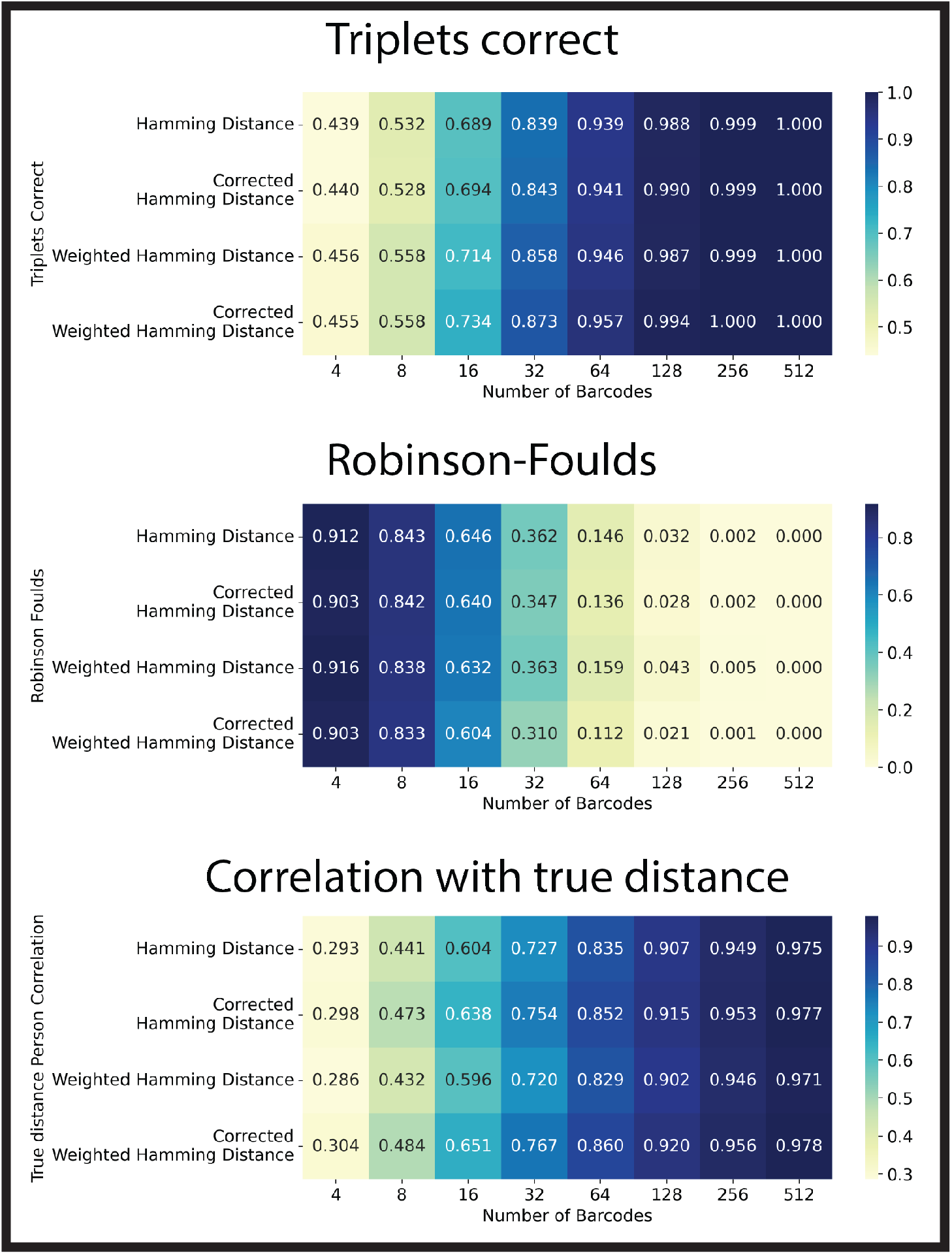
(Distance correction is consistent and more efficient than naive approaches) We bench-marked two baselines (NJ applied to Hamming Distance, as well as its weighted version) against the corrected versions which we propose. As the number of lineage tracing characters increased, the performance of all methods becomes perfect. This validates our theoretical consistency results for the distance correction scheme. Note that using the raw Hamming distance also provides consistent estimates of the trees. However, for a fixed number of lineage tracing characters, distance correction provides the best results. In other words, distance correction has a higher statistical efficiency compared to using the uncorrected Hamming distance. Each entry is the average performance over 50 repetitions.

